# HIV-producing resting T cells monopolize virus production, associated PANoptosis, and persistence in lymphoid tissue

**DOI:** 10.64898/2026.02.06.703840

**Authors:** Lijie Duan, Mark A. Sanders, Stephen W. Wietgrefe, Defeng Tian, Peter J. Southern, Jodi Anderson, Garritt Wieking, Timothy W. Schacker, Ashley T. Haase

## Abstract

Graphic:
In chronic HIV infection, HIV is produced before antiretroviral therapy (ART) by resting CD4^+^ T cells in lymphoid tissue germinal centers. Virus production is associated with destruction of T cells, follicular dendritic cells (FDCs), and B cells required for humoral immune responses by PANoptosis (apotosis, pyroptosis and necroptosis) mediated by the PANoptosome, a cellular structure that incorporates cell contents and programmed cell death pathways in a nodal latticework that mediates cell emptying and disruption in progressive stages indicated by the numbers. During ART, persistent HIV-producing resting T cells represent an immediate source of virus for infection rebound on treatment interruption.

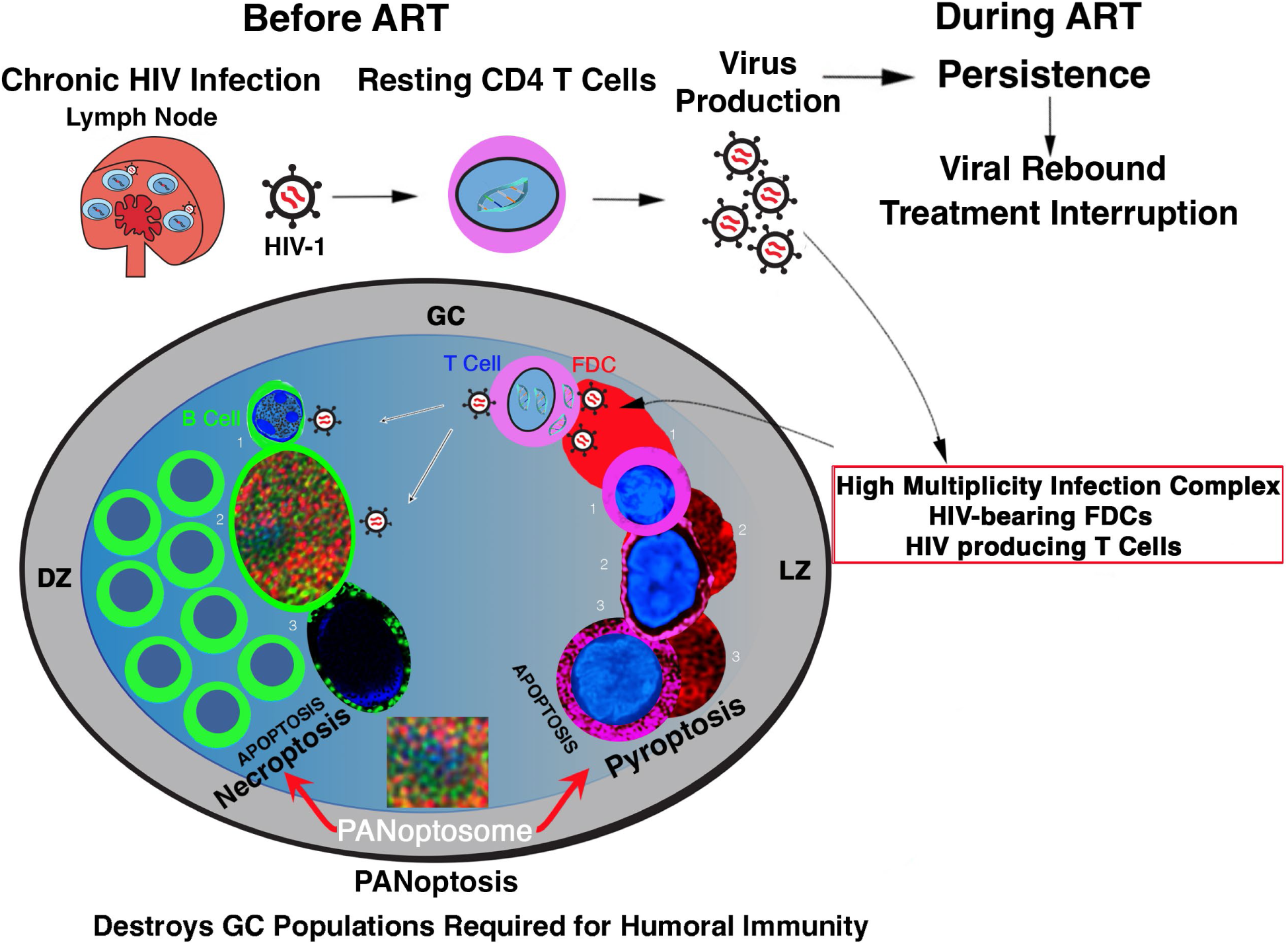

Activated CD4^+^ T cells have long been considered the principal source of HIV production before treatment and reactivated latently infected cells the source of virus rebound on treatment interruption, based largely on studies of peripheral blood T cells. Here we show before ART in the lymphoid tissue reservoir that HIV-producing resting CD4^+^ T cells are essentially the sole source of virus production in germinal centers. Virus is produced in infection complexes of virus bearing follicular dendritic cells (FDCs) that induce high multiplicity multi-HIV-DNA copy infections in overlying resting T cells. HIV production is associated with destruction by PANoptosis (pyroptosis, necroptosis and apoptosis) of interacting antibody-producing populations of FDCs, T cells and B cells. PANoptosis is mediated by the PANoptosome, visualized and defined here as a cellular structure incorporating cytoplasmic and nuclear contents and programmed cell death pathway components in a nodal latticework that executes cell emptying and disruption. During ART, HIV-producing resting T cells persist in lymphoid tissues when virus is undetectable in peripheral blood. These HIV-producing resting T cells that persist during ART represent an immediate source of virus to reignite infection on treatment interruption and are thus identified as an important new target for functional cure strategies.

## Introduction

HIV is generally thought to infect activated CD4^+^ T cells (1, 2) to give rise to productively infected and to latently infected resting T cell populations as a subset of the infected activated cells that returned to a resting state (3, 4). This latently infected cell population is a major focus of HIV cure research because recrudescent infection following interruption of ART is attributed to virus from reactivated latently infected cells (5–9). How this reconstruction of HIV production and persistence from analyses of *in vitro* cultured cells or peripheral blood T cells relates to virus production and persistence in the lymphoid tissue (LT) reservoir is unknown. Indeed, recent studies of the earliest detectable stage of HIV infection showed that HIV can directly infect resting T cells in lymphoid tissues to generate productively and latently infections populations, and the productively infected cells persisted for two to five years despite initiating ART within days of diagnosis (10). Thus, productively infected resting T cells could be an immediate source of virus for rebound following treatment interruption The potentially important implications of this possibility for devising and assessing new HIV cure strategies motivated the studies reported here of HIV infection of lymphoid resting T cells before and during ART, in the chronic stages of HIV infection when infection is usually diagnosed and ART initiated.

## Results

### Study participant lymphoid tissues

We analyzed archived lymph node (LN) biopsies available from a previously reported study (11) from participants who had been infected with Clade B HIV from 0.7 to 23 years. Plasma Viral Loads (pVL) ranged from 2530 to 157,000 copies per ml prior to initiating ART and were undetectable after 6 months of ART (Table 1).

**Table 1.** Participant ID (PID), demographics, duration of HIV Clade B infection and plasma viral load (VL) at Day 0 (D0) before and at month 6 of ART. (**11**).

| PID | Age | HIV infection duration<br>(years) | Plasma VL copies per ml |  |
| --- | --- | --- | --- | --- |
|  |  |  | D0 | ART Month 6 |
| 1950 | 23 | 4.6 | 2530 | <50 |
| 1954 | 24 | Unknown | 8370 | <50 |
| 1953 | 22 | 1.25 | 13,100 | <50 |
| 1860 | 30 | 0.7 | 141,000 | <50 |
| 1774 | 45 | Unknown | 157,000 | <50 |
| 1679 | 36 | 0.7 | 17,328 | <50 |
| 1669 | 23 | 23 | 6506 | <50 |

### Experimental Design

HIV-producing cells in LNs were identified and quantified using a direct and unambiguous assay for HIV-producing cells, defined as cells with visualized cell associated virions (12). The phenotypes of productively infected cells, cell-cell interactions, and cytopathology associated with virus production were determined by conventional histological, immunohistochemical and immunofluorescent techniques and microscopy for image capture and analysis. Lymphoid tissue sections were subsequently re-analyzed by super-resolution microscopy, using a Nikon Spatial Array Confocal (NSPARC) detector.

### Before ART, HIV is produced almost solely by CD4^+^CD25^-^ T cells in germinal centers

At D0 prior to initiating ART, nearly all (99%) of the HIV-producing cells were localized to the light cell zone (LZ) of germinal centers (GC) (Table 2; Figs. 1, 2A), and nearly all (99%) of the HIV-producing cells were fused and disrupted CD4^+^CD25^-^ Ki67^-^ T cells (Table 2; Figs. 1, 2B, 3).

**Fig. 1.**
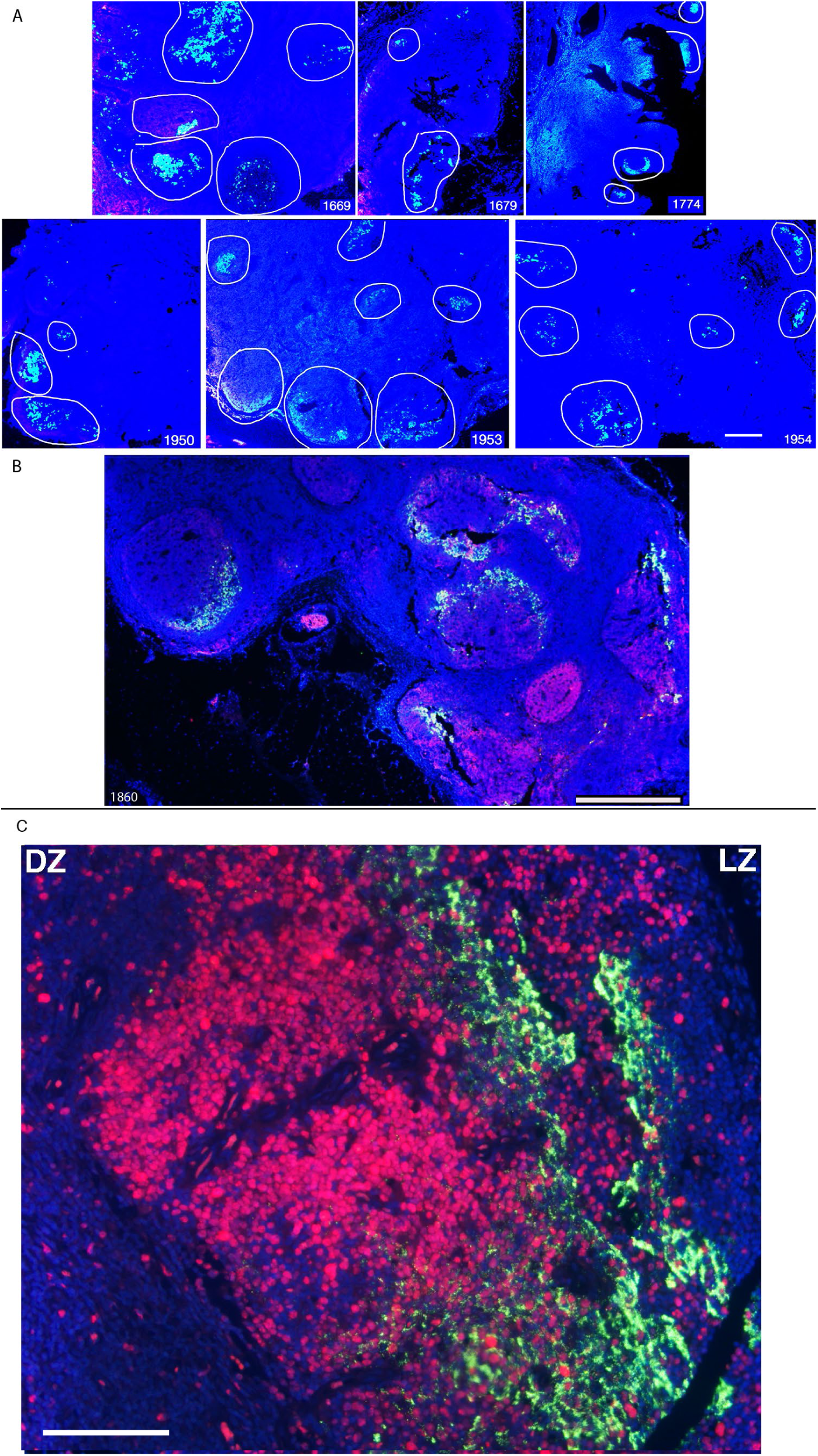
HIV-producing cells are concentrated in GCs in HIV infection. **A.** Each panel with patient ID (PID) in the lower right corner of the image shows a 4X view of blue green HIV-producing cell clusters in the germinal centers (GC) of B cell follicles delineated by white lines. Scale bar = 200 microns. **B, C.** HIV-producing cells and HIV RNA, blue green; CD20^+^ B cells, red. **C.** Ki67^+^ cells, red, to identify proliferating cells in the dark zone (DZ) of the GC. HIV-producing cells are Ki67^-^ cells in the light zone (LZ). Scale bar = 100 microns.

**Fig. 2.**
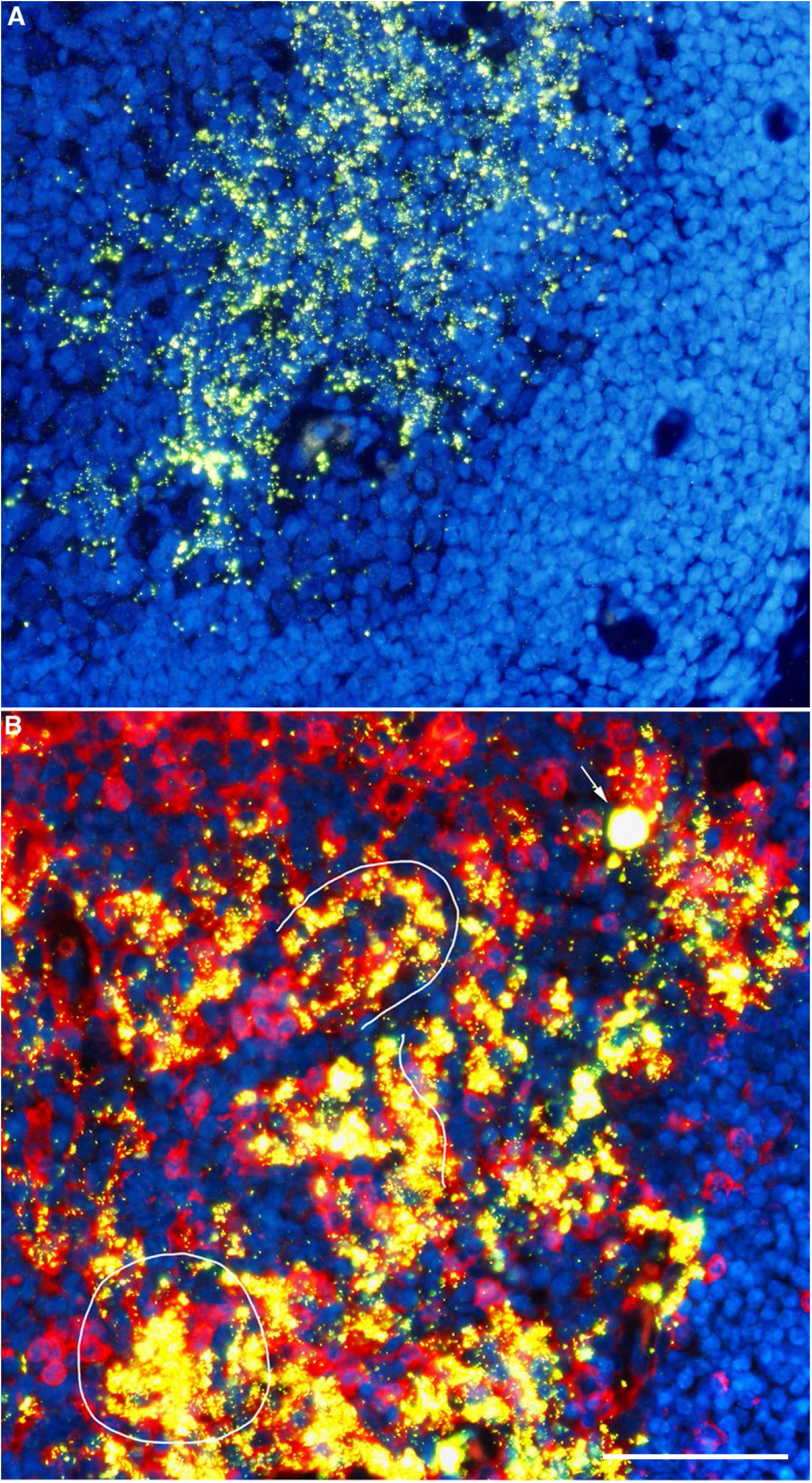
HIV-producing cells in GCs are fused and disrupted CD4^+^ T cells. Images representative of the LZs of the GCs shown in Fig.1. **A.** HIV RNA/virions, green; nuclei, blue. HIV virions in the LZ and HIV RNA and virions associated with fused and disrupted HIV-producing cells. **B.** HIV RNA/virions, green, yellow when associated with red CD4^+^ T cells; nuclei, blue. Image of GC at the junction with the follicular mantle zone (right border of the panel). Lines trace HIV cell-to-cell infection and fusion of adjacent CD4^+^ T cells and formation of multicellular aggregates (encircled). Most HIV virions are cell-associated in fused and disrupted cells (solid yellow white arrow points to an intact HIV producing CD4^+^ T cell). Scale bar = 100 microns.

**Fig. 3.**
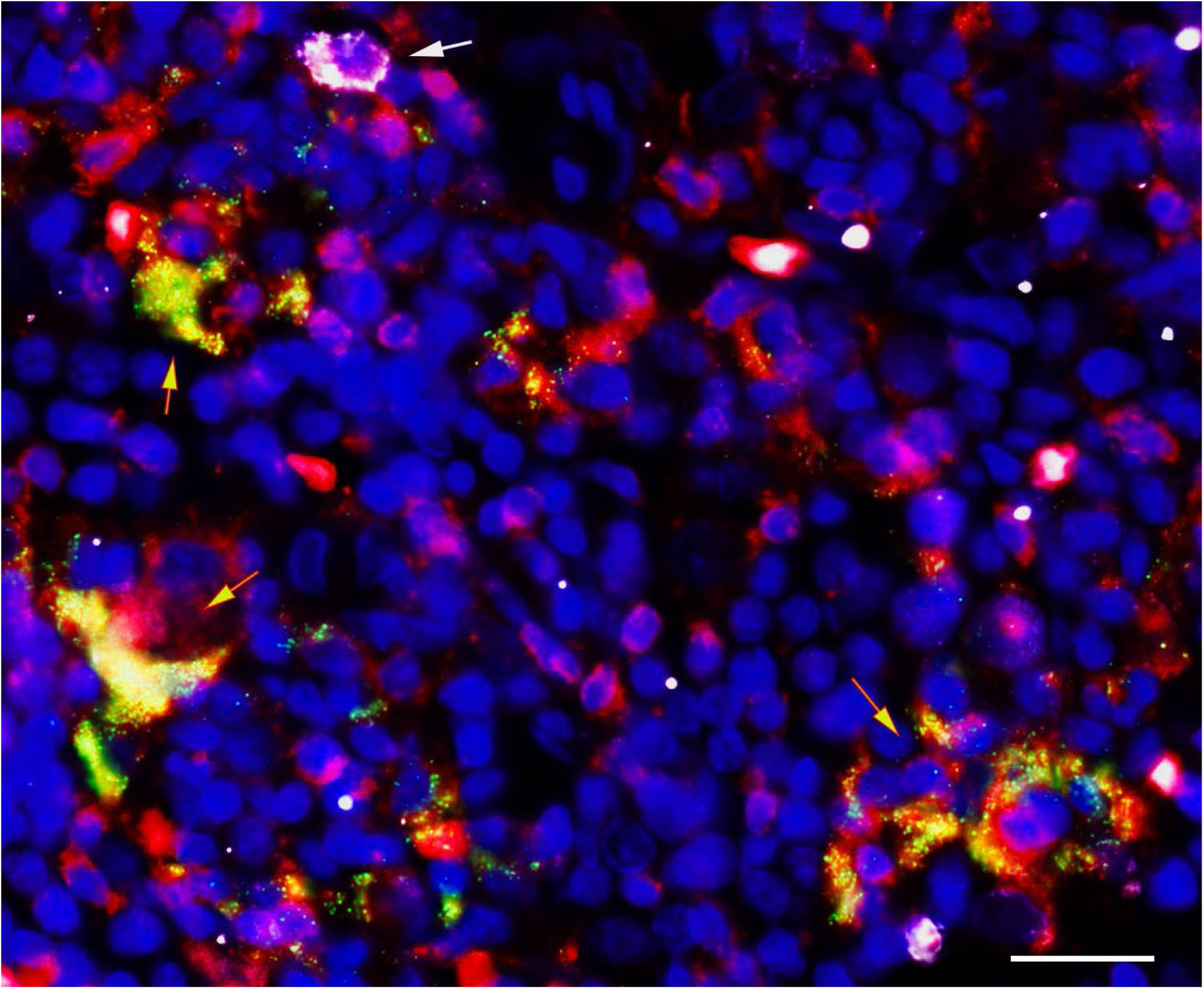
HIV-producing cells are spatially contiguous CD4^+^CD25^-^ T cells. HIV virions and RNA, green, yellow when associated with red CD4^+^ cells; CD25^+^ cells, white; nuclei, blue. Red-outlined yellow arrows point to cell-to-cell infection and fusion in multicellular HIV^+^ aggregates in spatially contiguous red CD4^+^ T cells. White arrow points to a HIV^-^CD4^+^CD25^+^ T cell. Scale bar = 50 microns.

**Table 2.**
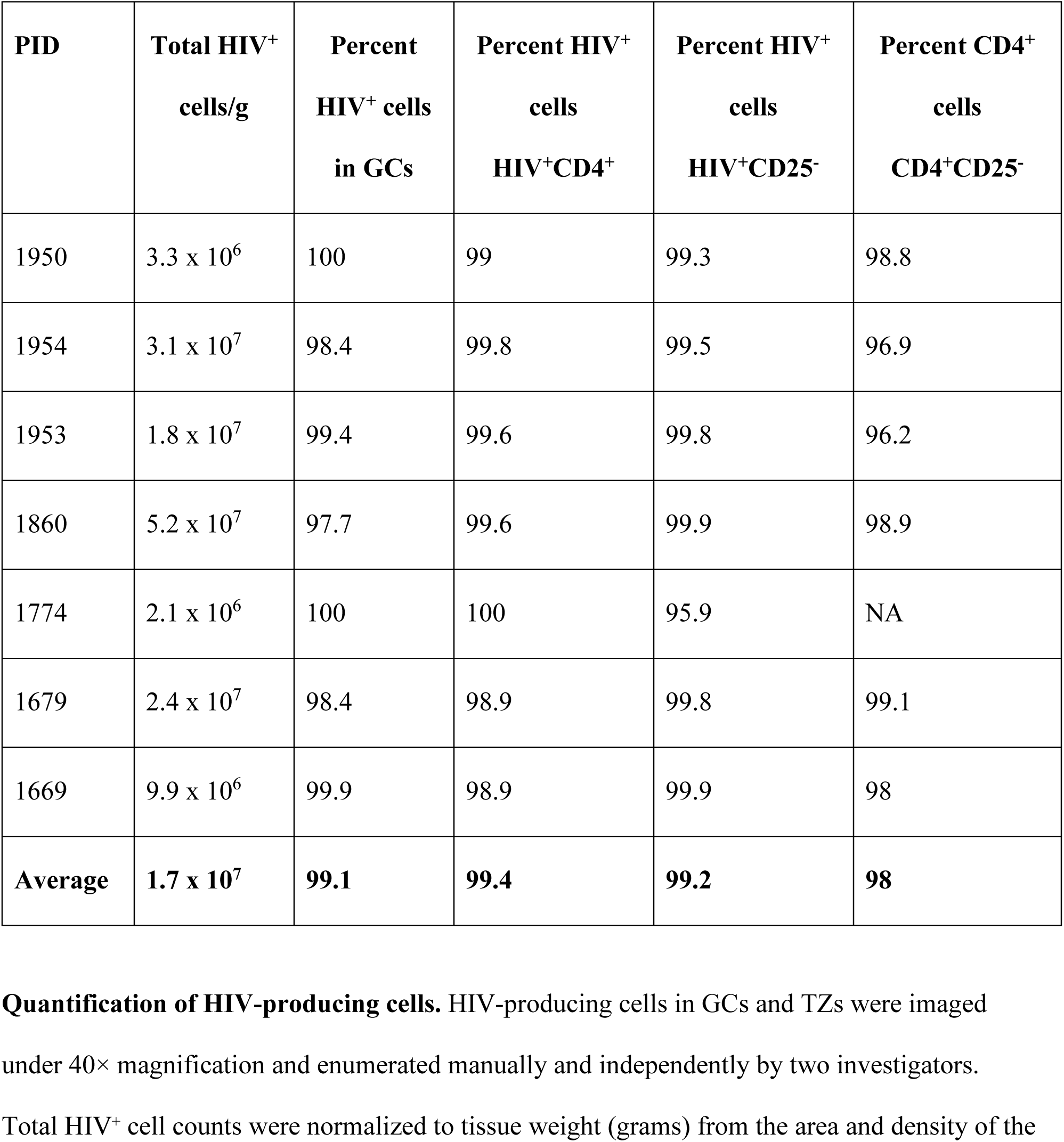
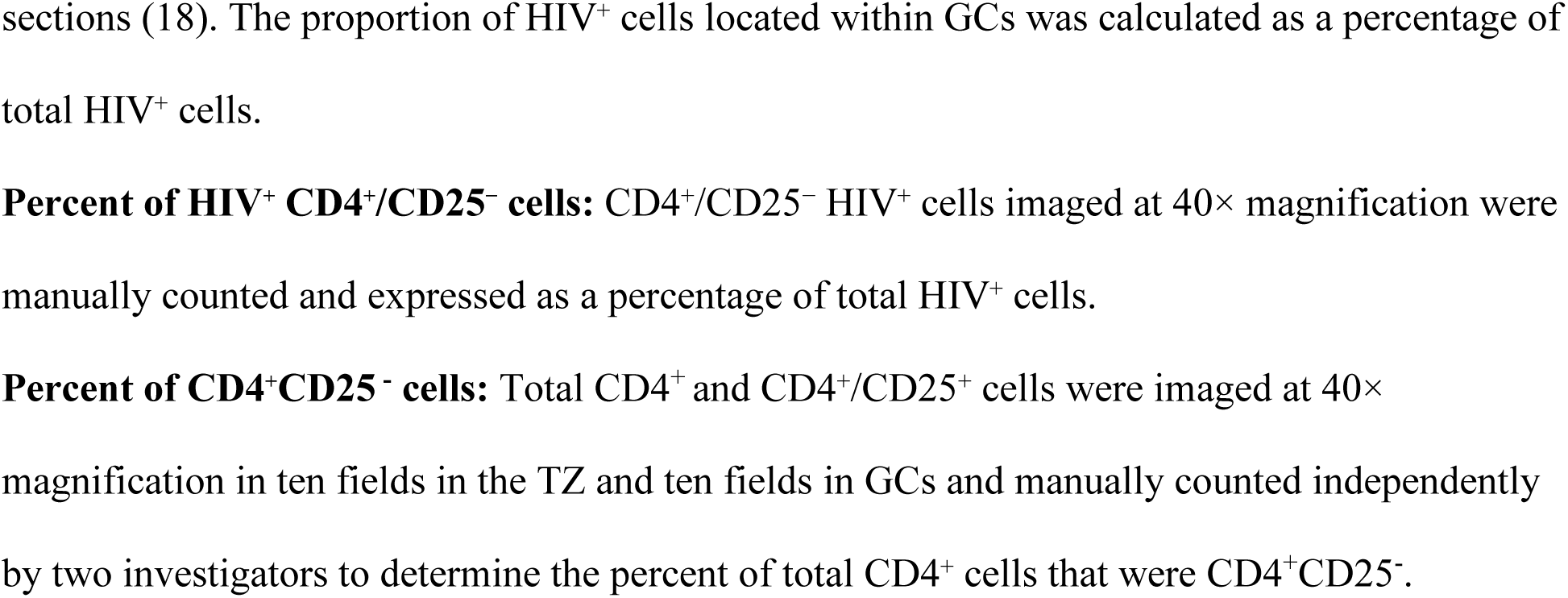
HIV-producing / CD4^+^CD25^-^ T cells in LN in chronic infection, D0, before ART.

### Availability and spatial contiguity concentrate infection of resting T cells in GCs

The nearly exclusive production of HIV by resting T cells in GCs was correlated with the predominance of susceptible resting T cell targets and their contiguity to each other and to virus associated with FDCs. HIV virus-producing cells were concentrated in the LZ with virions in a diffuse pattern typical of virus bound to FDCs (Fig. 2A), and the cells were fused together in a pattern reflecting HIV infection and fusion of adjacent CD4^+^ T cells to form multicellular aggregates in which most of the virus was cell-associated (Fig. 2B). Cell-to-cell infection and fusion of adjacent cells was localized to “islands” of CD4^+^CD25^-^ T cells (Fig. 3) in GCs in which 98 percent of the available CD4^+^ T cells in GCs were CD25^-^ (Table 2; Fig. 4). Beyond availability and spatial contiguity of resting T cell targets in GCs, infection was concentrated in GCs because the major available source of infection was virus bearing FDCs in a high multiplicity infection complex of FDCs fused to each other and to overlying resting T cells, as next described.

**Fig. 4.**
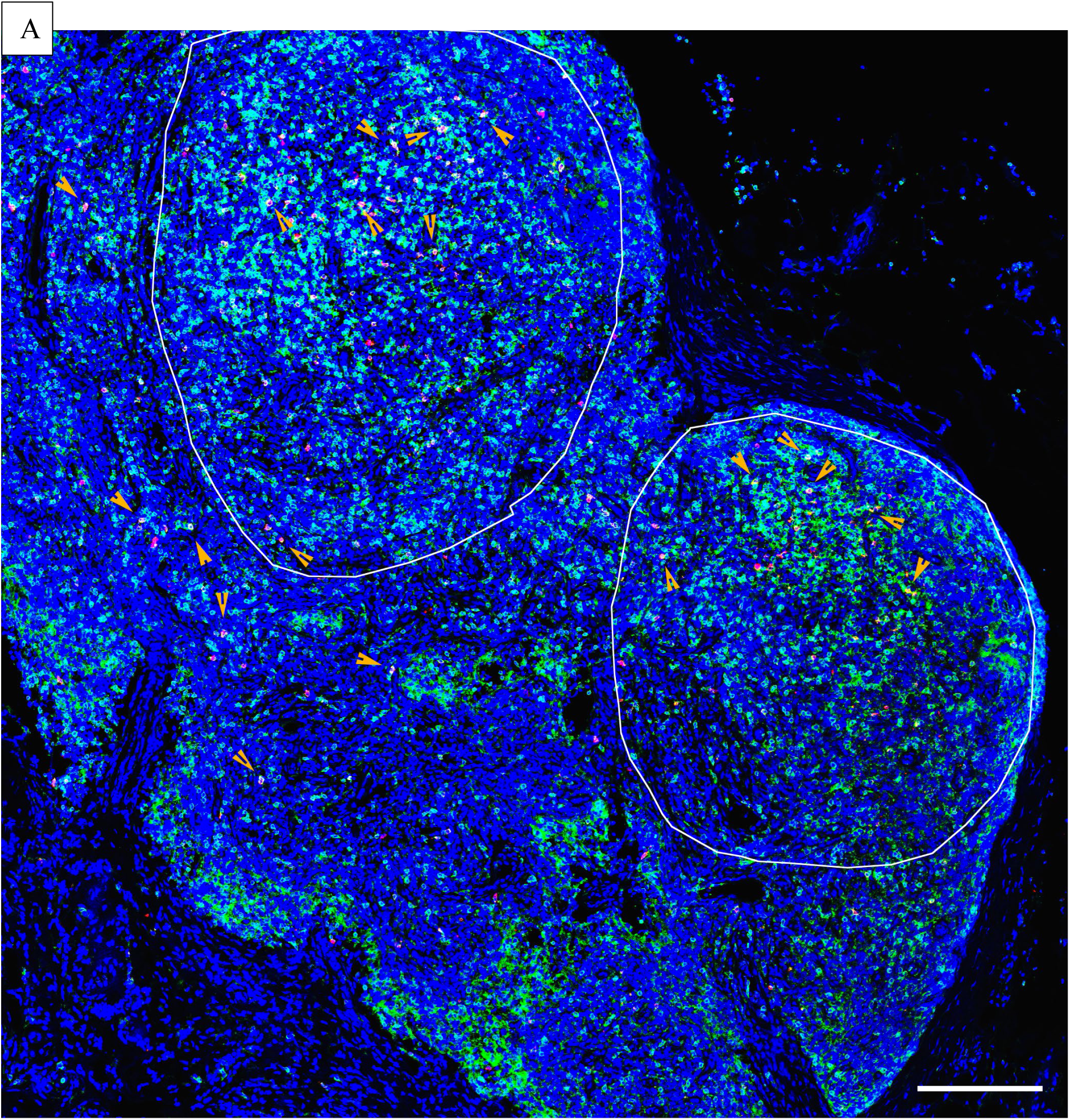
Predominant target cell availability of resting CD4^+^CD25^-^ T cells in GCs and TZ. The blue green cells in the encircled GCs and TZ are CD4^+^ T cells. Orange arrowheads point to the small number of CD4^+^CD25^+^ T cells. Of the HIV^+^ cells in GC, 99 percent were CD4^+^CD25^-^, closely corresponding the 98 percent of the HIV^-^CD4^+^ T cells that are CD25^-^ in GCs (Table 2). Scale bar = 200 microns.

### Visualizing HIV production in lymphoid tissues by super-resolution microscopy

The NSPARC detector for super-resolution microscopy enabled acquisition of images of HIV-producing cells in fixed LN tissue sections at <100 nm resolution with negligible background. These images confirmed the association of HIV virus and virus aggregates with fused and disrupted CD4^+^ T cells in GCs (Fig. 5). The mean diameter measured in 541,717 individual virions in 5 GCs was 138 nm, standard deviation, .008. This diameter is in good agreement with the expected average size of HIV virions of 120 nm (13) plus the added deposition of ELF-97 substrate for visualization.

**Fig. 5.**
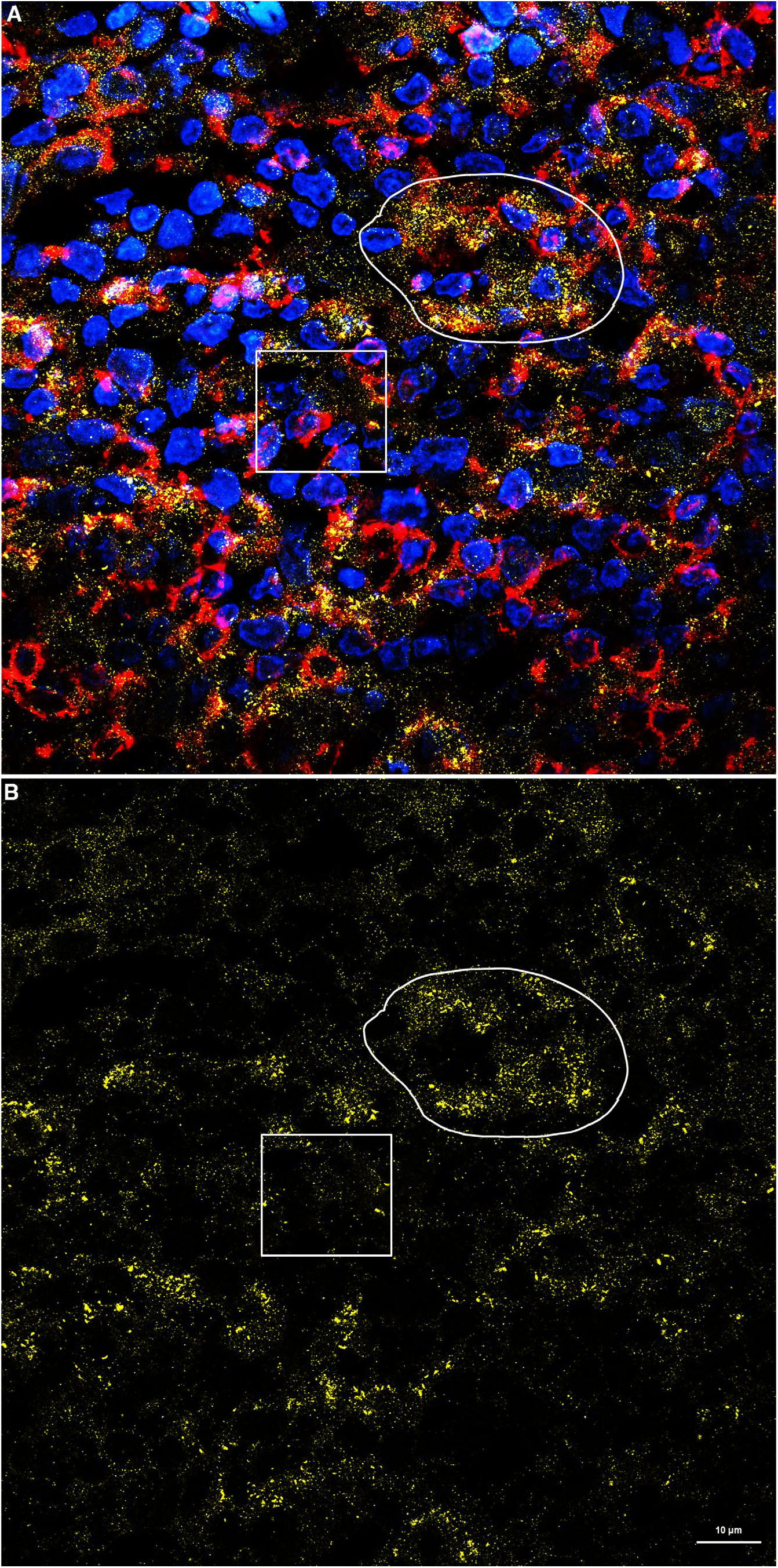
Super-resolution image of CD4^+^ T cell associated HIV in a GC. **A.** HIV, yellow; CD4, red; DAPI-stained nuclei, blue. **B.** HIV only. Scale bar, 10 µm. **A, B.** White traced region shows an example of HIV virions and virus aggregates mainly associated with fused and disrupted CD4^+^ T cells. The white box encloses HIV associated with nuclei of cells with no detectable CD4 expression from CD4^+^ T cells shown in subsequent Figures to be FDC-associated HIV. Scale bar=10 microns

### Infection fusion complex of CD4^+^ T cells and virus bearing FDCs

B cell follicles (BCF) and GCs have long been portrayed as engines of virus production where T cells migrating on the FDC network encounter virus, become infected and produce and deposit more virus on the FDCs to further amplify virus production (14–20). What the NSPARC super-resolution images of HIV-producing cells now revealed was an even more intimate relationship between virus source and targets in a high MOI (multiplicity of infection) fusion complex in which HIV-producing CD4^+^ T cells overlie and are fused to HIV-bearing FDCs (Overview Graphic and graphic in Fig. 6). We show images of this high MOI fusion complex in GCs in the coincidence of HIV virus and RNA (Fig. 6A), CD4^+^ T cells (Fig. 6B) and fused FDCs (Fig. 6C); and by the coincidence of CD4^+^ T cells and FDCs (Fig. 6B, C). The relationship between HIV replication in CD4^+^ T cells and HIV RNA^+^ FDCs is further supported by the colocalization of HIV virions and RNA in CD4^+^ T cells or portions thereof that superimpose on the portion of FDCs that are HIV RNA^+^ (Fig. 7).

**Fig. 6.**
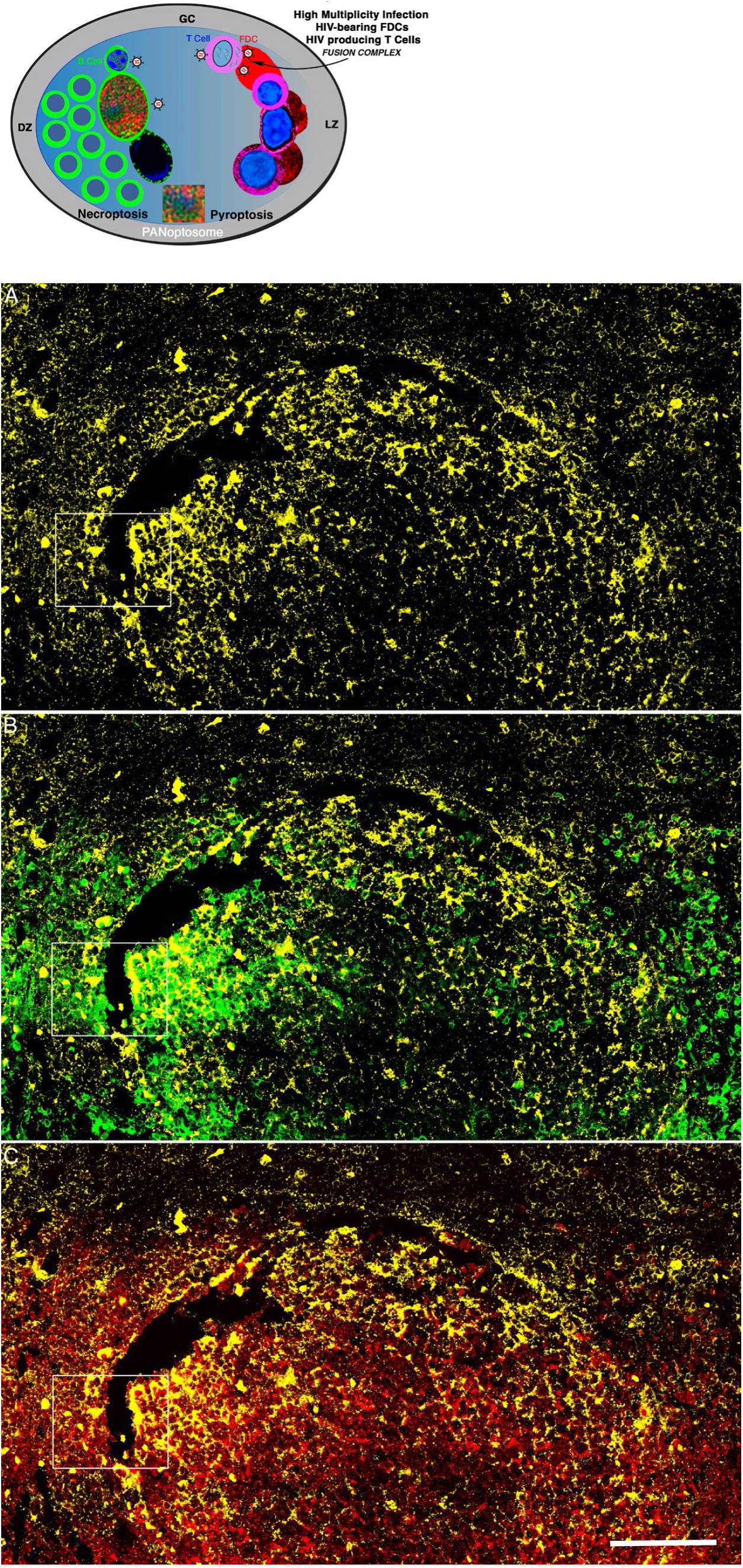
High multiplicity infectious fusion complex of HIV/CD4^+^ T cells and CD35^+^ FDCs. See graphic of T cells overlying HIV bearing FDCs. NSPARC Super-resolution images. HIV RNA/virions, yellow; CD4, green; CD35, red. HIV RNA and virions in the GC in **A** map onto HIV^+^CD4^+^ T cells shown in **B** and HIV^+^CD35^+^ FDCs shown in **C**; and the CD4^+^ T cells map onto FDCs (**B, C**). The white box encloses the same region in **A-C** shown in Fig. 7. Scale bar = 100 microns.

**Fig. 7.**
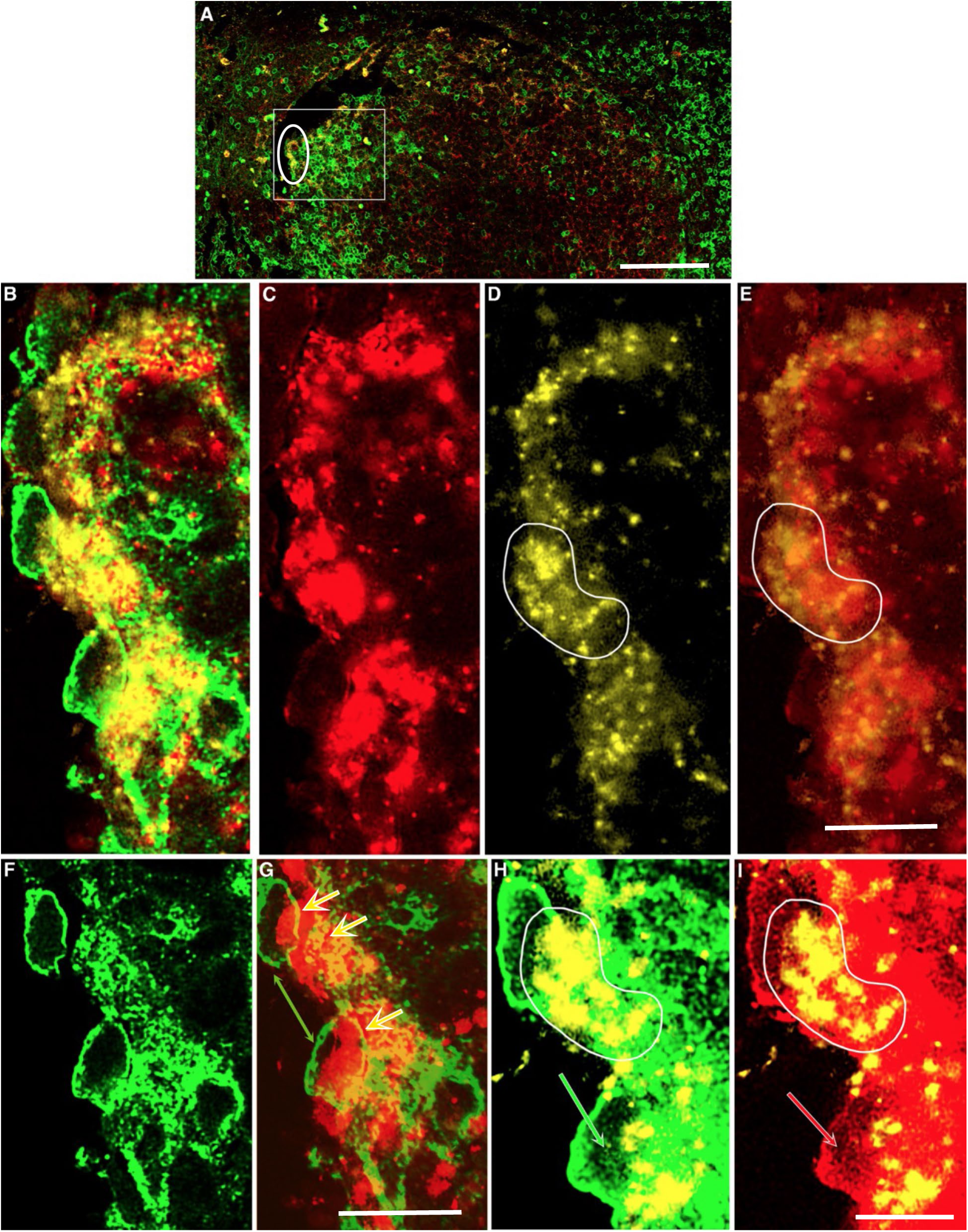
Offset superimposition of HIV in CD4^+^ T cells overlying HIV associated with FDCs. NSPARC super-resolution images. HIV^+^, yellow; CD35^+^ FDCs, red; CD4^+^ T cells, green. **A.** Scale bar = 100 microns. GC in Fig. 6 with box and circle enclosing images in **B-I.** Scale bar= 10 microns. **B.** HIV+ FDC + CD4. HIV virions, aggregates and RNA associated with fused and disrupted CD4^+^ T cells overlying HIV bearing fused FDCs. **C-E.** Scale bar = 10 microns. The FDC (**C**) and HIV (**D**) decreased opacity layers have been aligned to show the coincidence of HIV with FDCs in **E**. **F, G.** Scale bar = 10 microns. Aligned decreased opacity CD4 layer in F overlaid on FDC layer in **G**. **G**. Offset superimposition of two CD4^+^ T cells indicated by the green double-headed arrow. Yellow arrows point to fused CD4^+^ T cells or portions thereof fused with FDCs. **H, I.** The white traces in **D** and **E** enclose a region shown at higher magnification in **H** and **I.** Scale bar = 10 microns. HIV RNA colocalizes to FDCs, and the portion of the fused and disrupted CD4^+^ T cells or portions thereof superimposed on the HIV^+^ FDCs. White outlined green and red arrows point to nodal latticework in respectively partially emptied CD4^+^ T cells and FDCs.

### Multi-copy HIV DNA in resting memory T cells

High MOI fusion complexes of FDCs with multiple virions fused to overlying resting CD4^+^ T cells should induce multi-HIV-DNA copy infections. We tested this prediction by detecting HIV DNA in HIV-producing resting CD4^+^CD25^-^ T cells, which we additionally phenotyped as CD45RO^+^ memory cells. We did detect cells with the single copy of HIV DNA in their nuclei expected for infections initiated with a single virion in cells spatially separated from the infection complexes of T cells and FDCs. However, we detected far more abundantly the predicted large numbers of copies of HIV DNA in GCs in a pattern resembling the predicted high MOI infections in T cell/FDC infection complexes (Fig. 8). In the NSPARC super-resolution images, we documented multiple copies and aggregates of HIV DNA in the disrupted cytoplasm and nuclei of multinucleated CD45RO^+^ memory cells; in cells with apoptotic appearing nuclei; and in small memory cells mainly in the cytoplasm (Fig. 9). These images are consistent with a reconstruction of the *in vivo* HIV life cycle in the high MOI FDC-T cell fusion complex of ongoing synthesis of HIV DNA in the cytoplasm followed by nuclear location in multiply infected T cells.

**Fig. 8.**
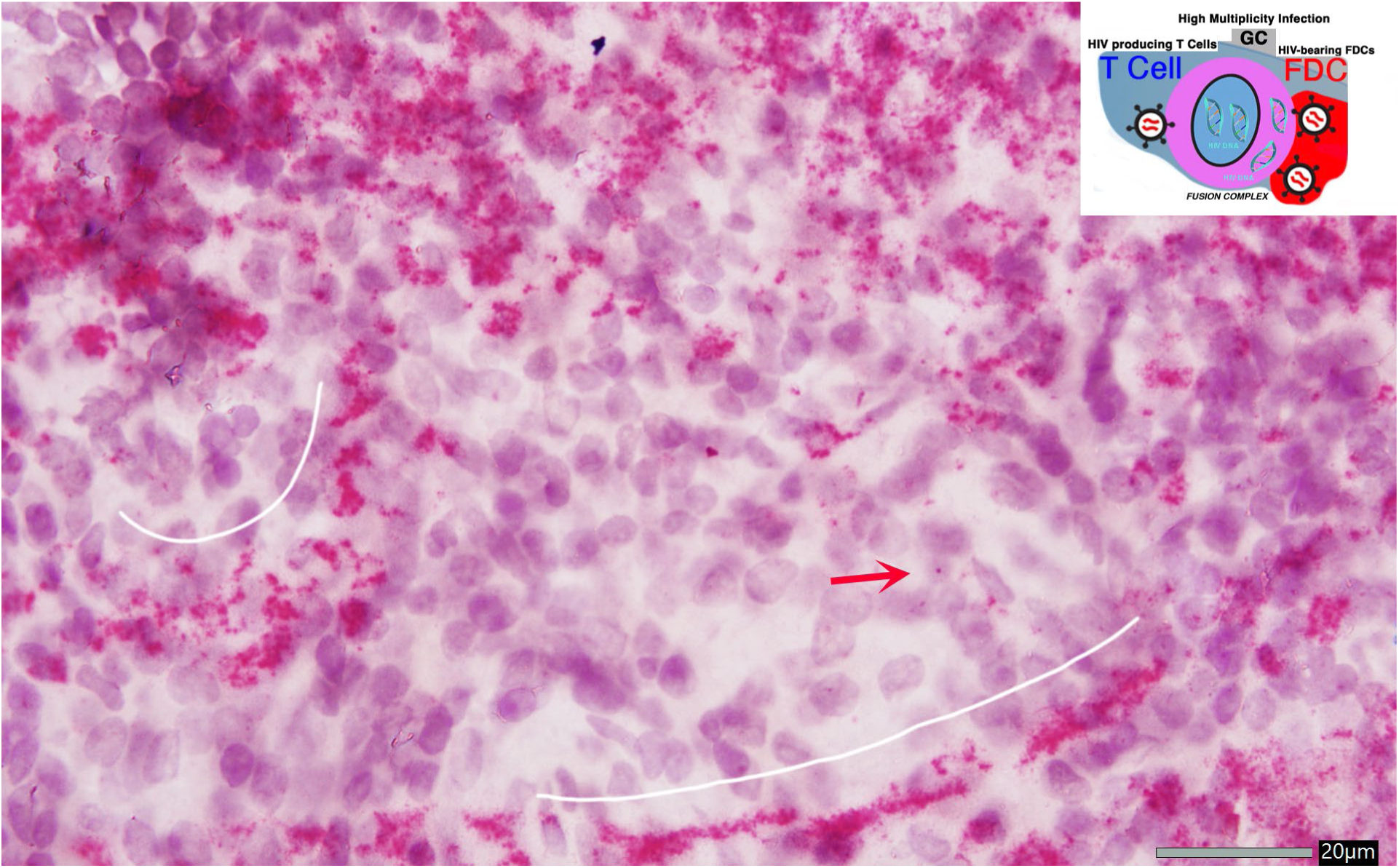
Multiple copies of HIV DNA produced in a high multiplicity infection complex of HIV bearing FDCs fused to resting memory T cells. Inset from the Graphic illustrating high MOI infection of T cells by HIV bearing FDCs that predicts multi-DNA copy infections. HIV DNA, red. White lines trace multiple copies of HIV DNA in fused cells in the LZ of adjacent GCs. Note the single copy of HIV DNA (red arrow) in the nucleus of a cell inside of the infection complex. Scale bar =20 microns.

**Fig. 9.**
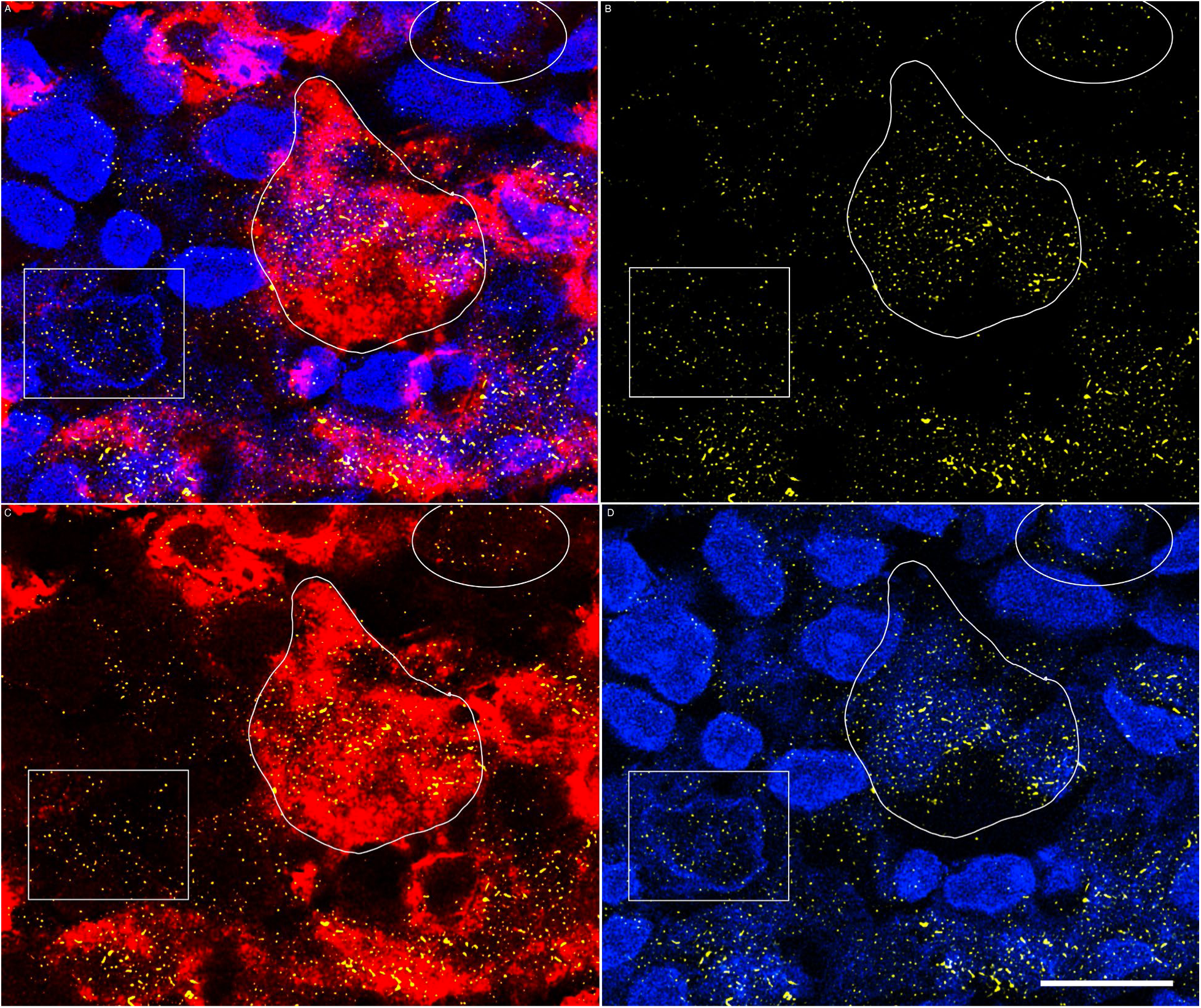
Multiple copies of HIV DNA in memory T cells. NSPARC super-resolution images. HIV DNA, yellow; CD45RO^+^ memory cells, red; DAPI-stained nuclei, blue. **A.** HIV DNA + CD45RO + DAPI-stained nuclei. **B.** HIV DNA. **C.** HIV DNA + CD45RO. **D.** HIV DNA + DAPI-stained nuclei. White traces in **A-D** enclose a multinucleated CD45RO^+^ cell with multiple punctate copies and aggregates of HIV DNA (**B**) associated with the cytoplasm (**C**) and nuclei (**D**) of the cells. The white boxes enclose cells with HIV DNA (**A, B**) associated with cytoplasmic remnants (**C**) and nucleus of the cell with apoptotic changes of margination of the nuclear envelope and nuclear fragments (**D**). The circle encloses a small, disrupted cell with HIV DNA mainly in the cytoplasm (**A-D**). Scale bar = 10 microns.

### Pyroptosis of FDCs and PANoptosis of T cells

In investigating the striking cytopathic effects associated with virus production of fusion, syncytium formation, cell disruption and destruction of T cells and FDCs, we found that the cells were succumbing to pyroptosis and PANoptosis. PANoptosis generally refers to pyroptosis, necroptosis and apoptosis in cell populations and also to individual cells that manifest one or more of the three major forms of programmed cell death (PCD) (21–23). Here, PANoptosis refers to pyroptosis and apoptosis of CD4^+^ T cells and the PANoptosome to a previously documented cell structure in type II pneumocytes in COVID-19 pneumonia (23) that empties and disrupts cells through a nodal latticework. We show images (Fig. 10A-D) of FDCs undergoing pyroptosis and overlying HIV-producing cells undergoing pyroptosis + apoptosis with a graphic (Fig. 10, right) that provides the following interpretative framework: 1) asynchronous cell death processes in tissues enable visualization of stages of the process in a single snapshot in time; 2) illustrations of pyroptotic changes in the cytoplasm of T cells and FDCs and nuclei of T cells in PANoptosis (apoptosis and pyroptosis) simplified to three stages; and 3) definition of the PANoptosome. In the actual images (Fig. 11) of the cytoplasm and nuclei of pyroptotic/PANoptotic (pyroptosis+apoptosis) CD4^+^ T cells, stage 1 cells have relatively normal staining of cytoplasm and nucleus. In stage 2, the cytoplasmic rim of CD4 is thinned and disrupted and the nuclei show apoptotic changes of margination of the nuclear envelope and fragments. In stage 3, only the nodal latticework associated with emptying of cell contents is visible. We further document PANoptosis (pyroptosis + apoptosis) of CD4^+^ T cells by overlaying images of Caspase-1 (pyroptosis), Caspase-3 (apoptosis) and HIV RNA, and found that most of the Caspase-1^+^/Caspase-3^+^ PANoptotic T cell fusions with pyroptotic Caspase-1^+^ FDCs included both infected and uninfected T cells (Fig. 12). In the super-resolution images of pyroptosis of the T cell-FDC complex (Fig. 13), the syncytium of CD4^+^ T cells and FDCs in GCs are comprised of largely emptied stage 3 cells intermixed with stage 1 multicellular fusions of HIV^+^CD4^+^ T cells or HIV^+^CD35^+^ FDCs (Fig. 13A). A high magnification view of two superimposed CD4^+^ T cells and FDCs shows the residual latticework nodes at the rim of the largely emptied CD4^+^ T cells and the nodes and arms of the latticework in the underlying FDCs (Fig. 13B-D). The coincidence in HIV (and by inference, CD4) and Gasdermin D (GSD) (for pyroptosis) in nodes in fused CD35^+^ FDCs satisfies the definition of the PANoptosome as a nodal structure that incorporates cellular, in this case, viral, and pyroptotic cell death pathway components in the nodes of the cytoplasm of pyroptotic fused FDCs (Fig. 14; Fig. 15A-D) and nuclear contents incorporated into a nodal latticework (Fig. 15E). The movie (Supplemental Fig. 1) shows the nodes and their contents distributed throughout the cytoplasm of the fused FDCs in 3D and also highlights the asynchrony of their emptying in the z slice down comparison of smaller nodes in a more advanced stage of emptying with larger nodes at an earlier stage.

**Fig. 10.**
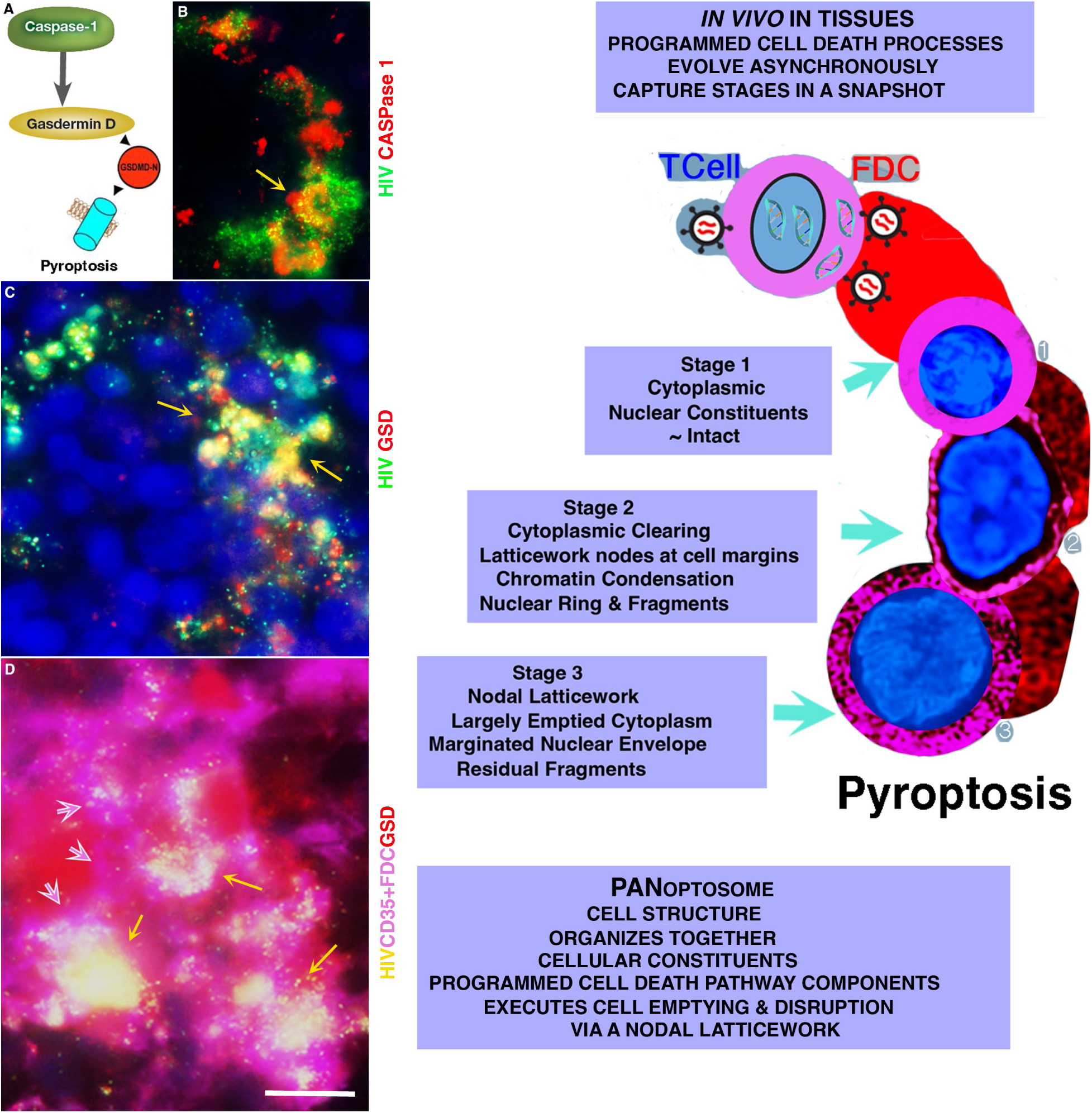
Pyroptosis of HIV-producing CD4^+^ T cells and FDCs and graphic of stages of cell clearing and disruption mediated by the PANoptosome. Left panels: **A.** Cartoon of the pyroptotic pathway: Caspase-1 cleavage of Gasdermin D to generate pore forming N terminal GSDMD-N (abbreviated, GSD) that mediates pyroptosis. **B.** HIV RNA, green; Caspase-1, red. GC in which fused and disrupted FDCs are Caspase-1^+^; yellow arrow points to Caspase-1^+^ T cells overlying FDCs. **C.** HIV, green; GSD, red. GC in which yellow arrows point to HIV^+^GSD^+^ cell fragments in pyroptotic T cells overlying red pyroptotic fused and disrupted FDCs. **D.** Red GSD^+^ fused magenta CD35^+^ FDCs (border of one cell marked by white outlined magenta arrows) with HIV^+^GSD^+^ fused cells (yellow arrows) overlying FDCs. Scale bar =10 microns. Right Graphic: Introduction to pyroptosis of the T cell/FDC fusion complex and PANoptosis and the PANoptosome. Illustration of three numbered stages of pyroptosis of T cells and FDCs in NSPARC super-resolution snapshots of an asynchronous process of cell clearing and disruption executed by the PANoptosome nodal latticework. PANoptosome defined in the graphic.

**Fig. 11.**
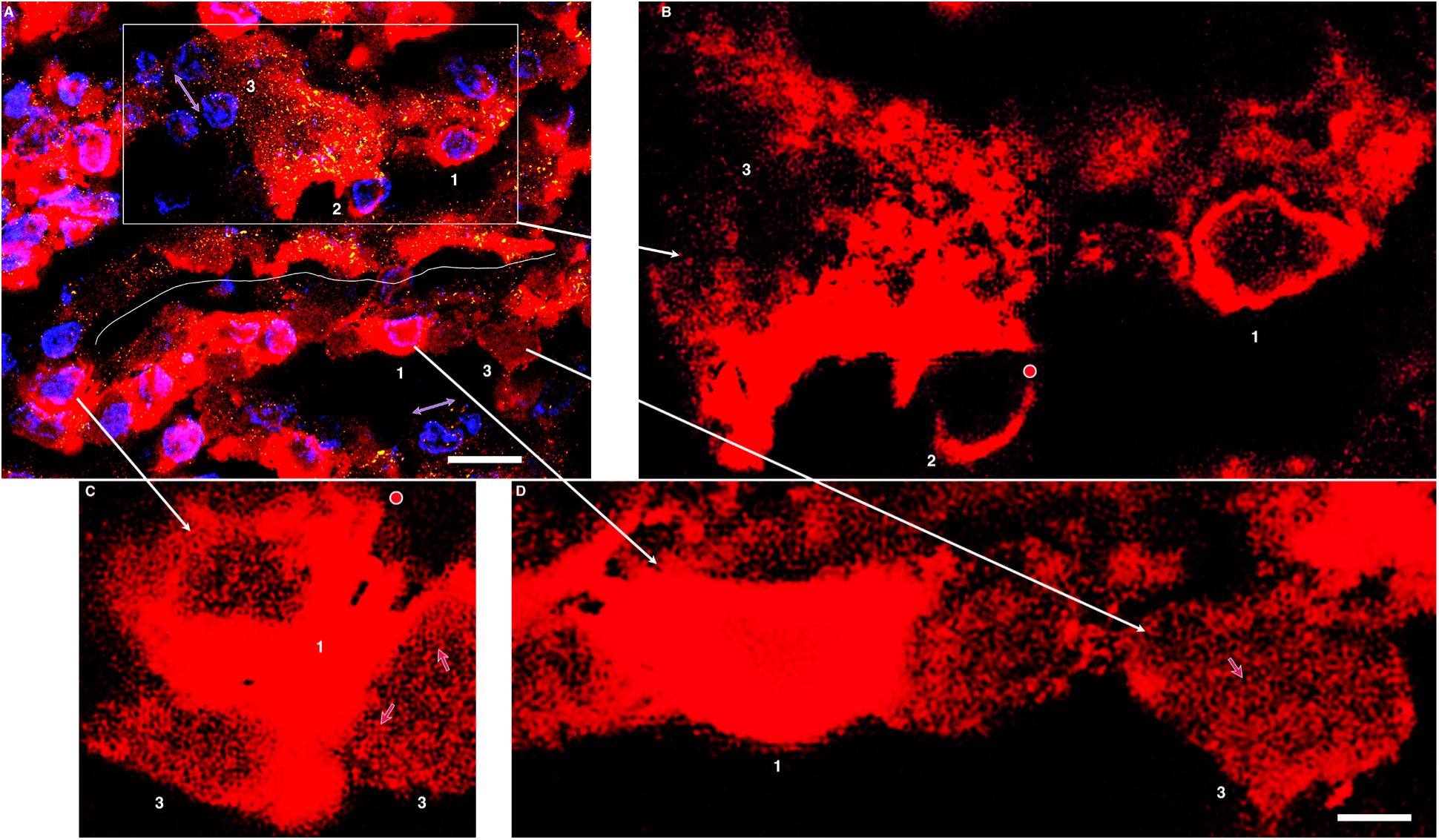
Pyroptosis of CD4^+^ T cells and cell emptying and disruption via a nodal latticework. NSPARC super-resolution images. HIV, yellow; CD4, red; DAPI-stained nuclei, blue. **A.** HIV+CD4+DAPI. Scale bar = 20 microns; **B-D**, CD4. **A.** White box and line enclose respectively a multicellular fusion complex of CD4^+^ T cells with associated virions and virus aggregates and cell-to cell infection and fusion. Numbers refer to stages of cell emptying described in the Fig. 10 graphic. Lavender double-headed arrows point to stage 2 and 3 nuclear changes of envelope margination and fragments. White arrows connect cells to enlarged images in **B-D.** Scale bar = 5 microns. **B.** Enlarged view of stages of cell emptying; numbers of stages as in **A**. Encircled red dot is adjacent to nodes in the thinned CD4^+^ cytoplasmic rim in the stage 2 cell. **C.** Fusion complex of stage 1 and 3 cells. Red dot is adjacent to nodes and latticework at the margins of the emptying stage 1 cells. Arrows point to latticework and arms and nodes of the latticework. **D.** Comparison of stage 1 and 3. Numbers and arrow as described in **A-C**.

**Fig. 12.**
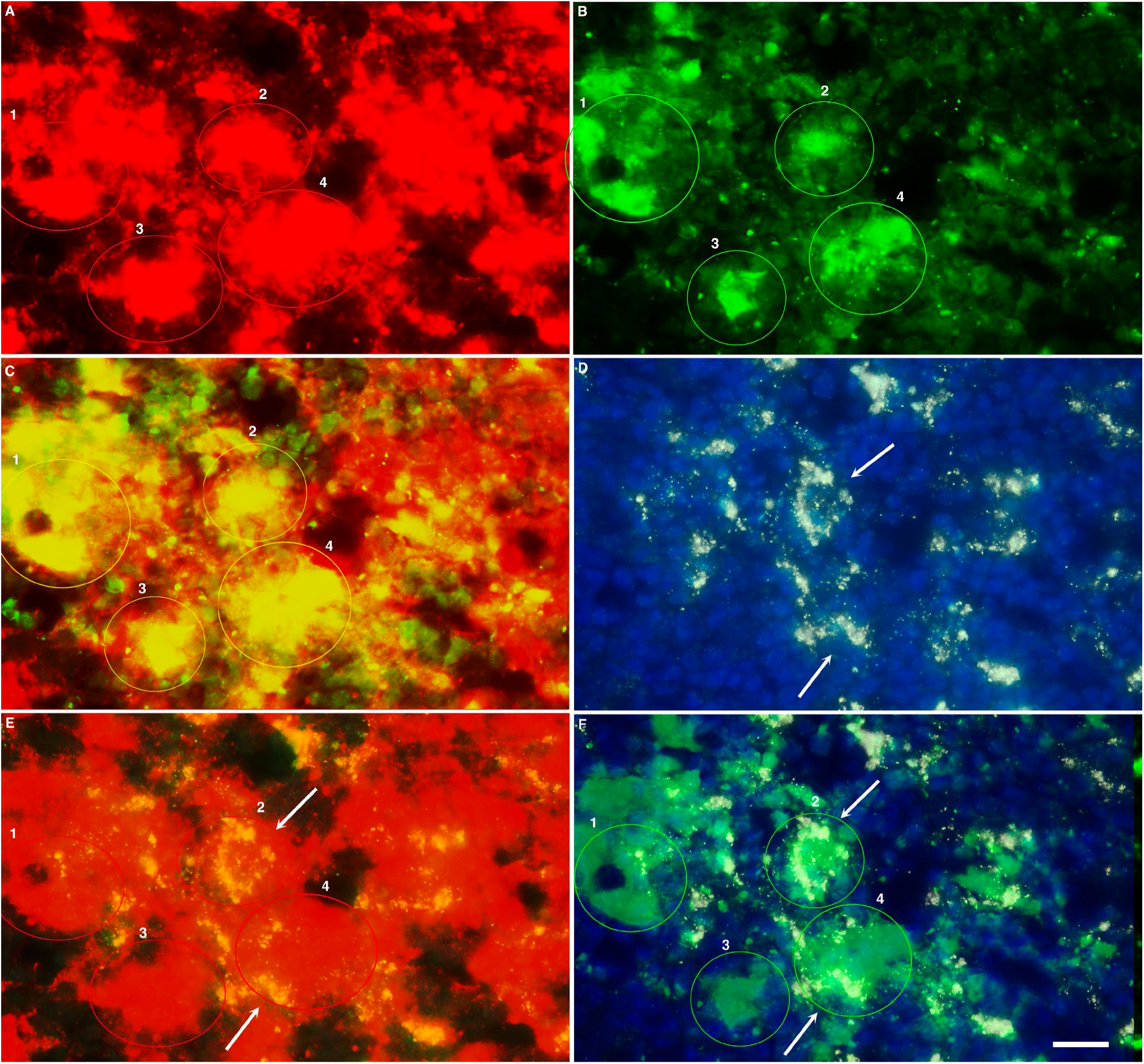
PANoptosis (Pyroptosis + Apoptosis) of HIV^+^ and HIV^-^ cells and pyroptosis of FDCs. Caspase-1, marker for pyroptosis, red; caspase-3, marker for apoptosis, green; HIV, white; DAPI-stained nuclei, blue. **A-C. A.** Caspase-1^+^ fused pyroptotic FDCs. Four numbered multicellular fusions (red circles) correspond to caspase-3^+^ fused cells in **B** (green circles) and the yellow encircled caspase-1^+^caspase-3^+^ pyroptotic/apoptotic fused T cells overlying pyroptotic FDCs in **C**. **D-F. D.** HIV. White arrows point to fused and disrupted HIV^+^ cells. **E, F.** Decreased opacity overlaid images of caspase-1, HIV and caspase-3. White arrows in the encircled fused cells point to HIV^+^ PANoptotic (pyroptosis+apoptosis) yellow HIV^+^ Caspase-1^+^ cells in **E** and green /white HIV^+^Caspase-3^+^ cells in **F**. Scale bar = 20 microns. Most of the PANoptotic fused cells are HIV^-^.

**Fig. 13.**
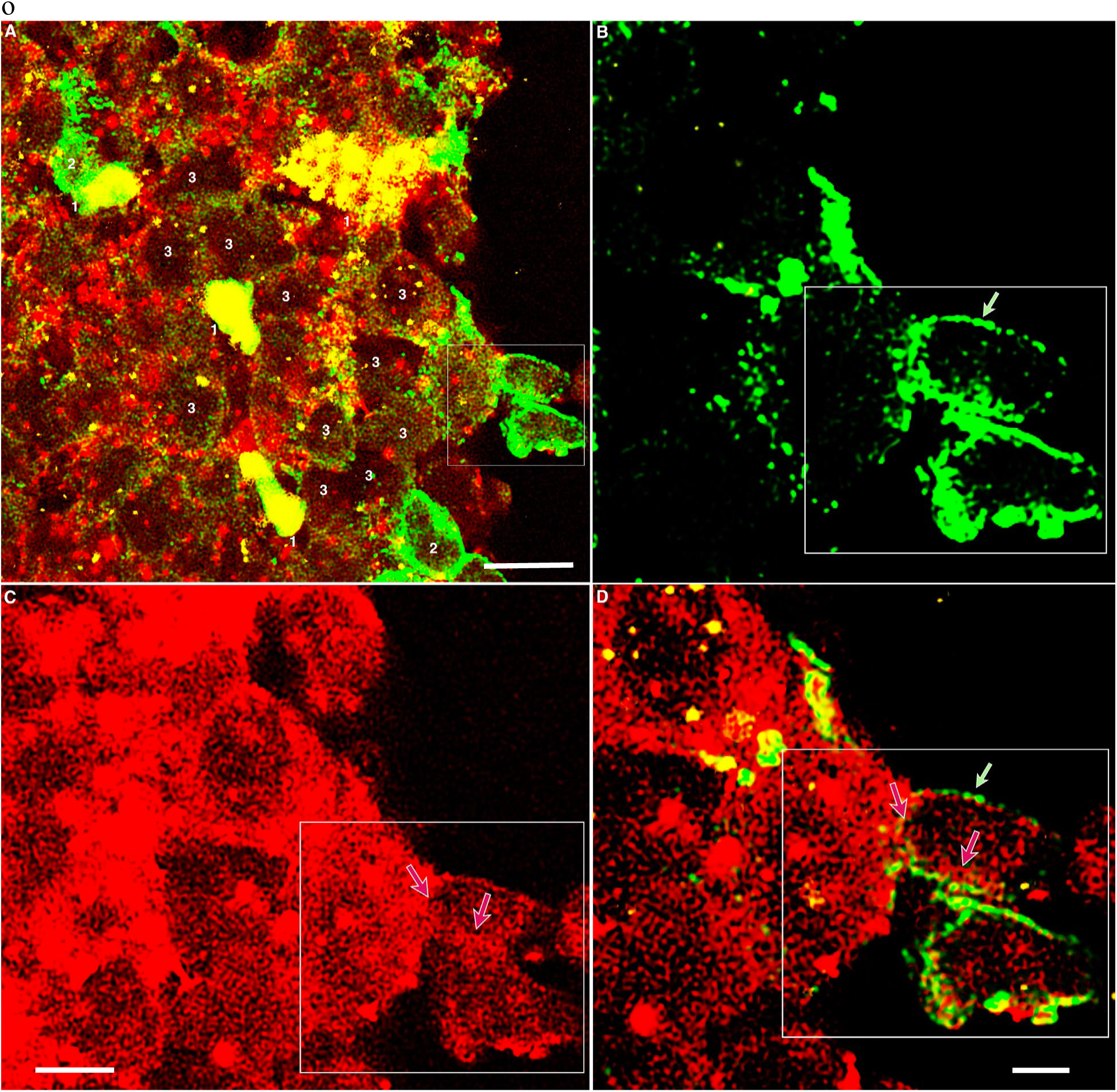
Pyroptosis of the fused infection complex of HIV producing CD4^+^ T cells and FDCs with cell emptying and disruption via a nodal latticework. NSPARC super-resolution images. HIV, yellow; CD4^+^ T cells, green; CD35^+^ FDCs, red. **A.** HIV+CD4+CD35; white box encloses two CD4^+^ T cells superimposed on two CD35^+^ FDCs. Scale bar = 10 microns. Higher magnification of CD4 in **B**, CD35 in **C** and CD4+CD35 in **D**. Numbers refer to stages of cell emptying described in the Fig. 10 graphic. **A.** Syncytium of mainly stage 3 fused T cells and FDCs. Cells marked stage 1 are multicellular fusions of HIV^+^CD4^+^ T cells or HIV^+^CD35^+^ FDCs. Two cells are marked as examples of stage 2. **B.** Outlined green arrow points to CD4^+^ nodes at the cell margins in largely emptied cells. **C.** Scale bar =5 microns. Outlined red arrows point to the nodes and arms of the FDC latticework. **D.** Scale bar = 5 microns. Superimposed imprint of CD4^+^ T cells fused to FDC nodal latticework. Arrows as described in **B** and **C**.

**Fig. 14.**
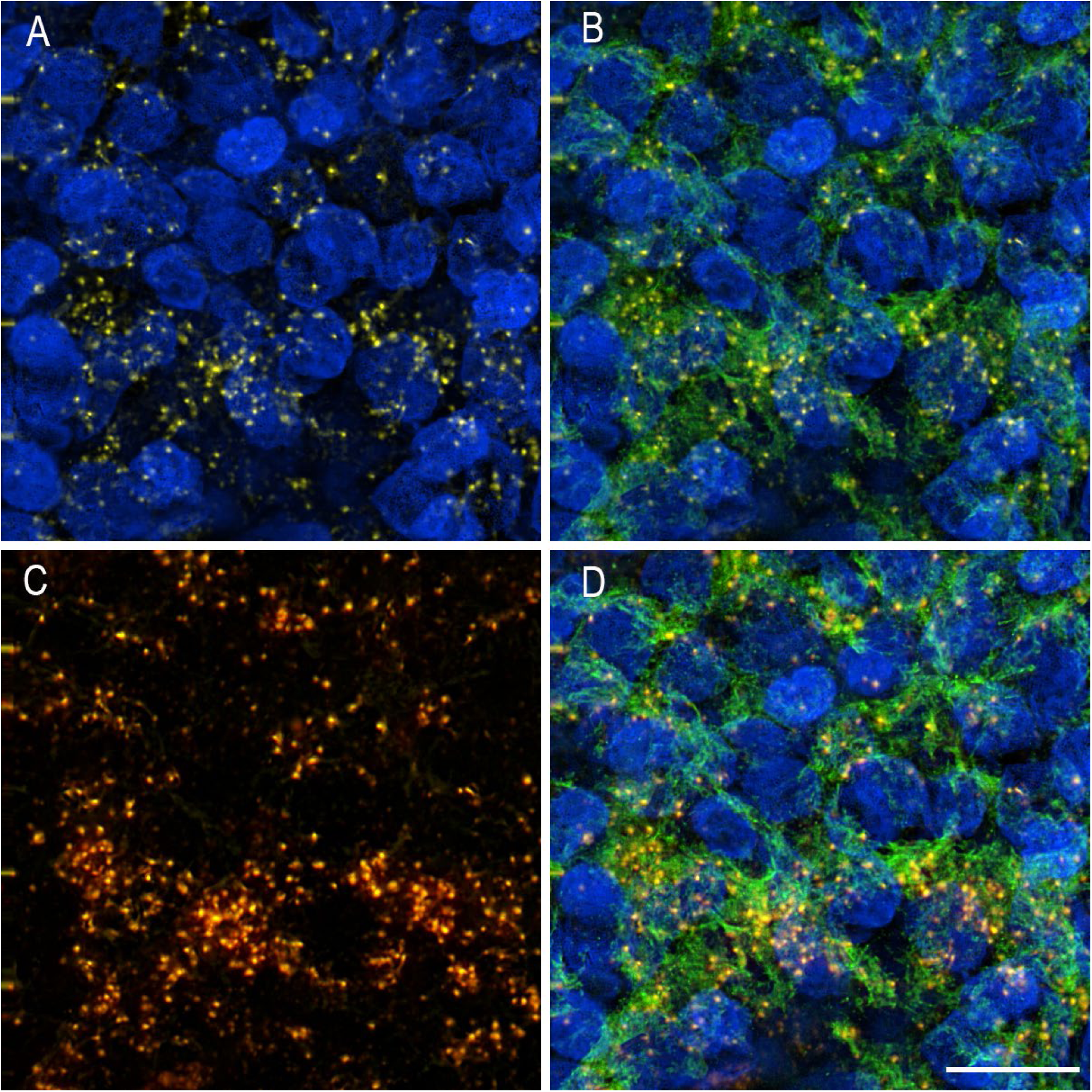
HIV^+^ GSD^+^ nodes in pyroptotic CD35^+^ FDCs. NSPARC Super-resolution images. HIV, yellow; CD35+, green; GSD, red; nuclei, blue. **A.** HIV^+^ nodes. **B.** HIV^+^ nodes in fused CD35^+^ FDCs. **C.** HIV^+^GSD^+^ nodes. **D.** Coincidence of HIV^+^GSD^+^ nodes in CD35^+^ FDCs. Scale bar = 10 microns.

**Fig. 15.**
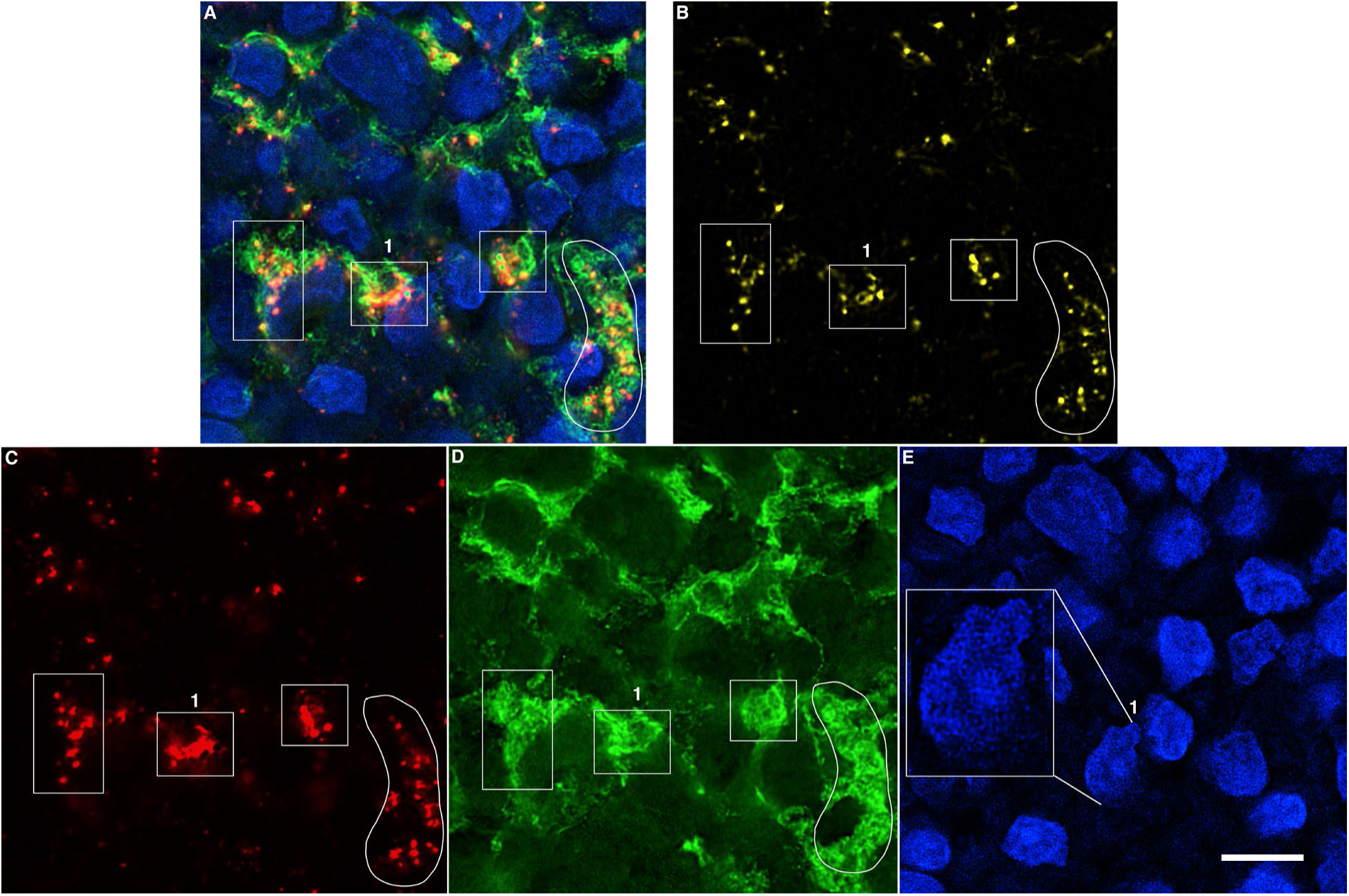
HIV^+^CD35^+^ GSD^+^ nodes and nuclear latticework in pyroptotic FDCs. NSPARC Super-resolution images. HIV, yellow; CD35+, green; GSD, red; Dapi-stained nuclei, blue. **A.** HIV+CD35+GSD+Dapi. B. HIV. C. GSD. D. CD35. E. Dapi-stained nuclei. **A-D**. White traces and boxes enclose examples of the coincidence of HIV, GSD and CD35 in yellow, green and red orange nodes. **E.** Exploded view of nuclear nodal latticework in the boxes labelled 1. Scale bar = 10 microns.

### Necroptosis and PANoptosis of B cells

We also discovered that B cells were undergoing necroptosis and PANoptosis (apoptosis and necroptosis) at the interface between the HIV-producing T cells and FDCs in the LZ. We provide a graphic showing HIV produced by T cells binding to B cells without replication and three stages of emptying mediated by the PANoptosome as an interpretative framework for the super-resolution image analysis (Fig. 16) that shows (Figs. 17, 18) HIV virions bound to CD20^+^ B cells without evidence of replication (i.e., intra-cellular HIV RNA^-^) in a syncytium of fused cells undergoing necroptosis associated with a CD20^+^RIPK3^+^ nodal latticework (Fig. 17 A-D). Cells characterized as stage 1 are small with a largely intact rim of CD20^+^ cytoplasm and low levels of RIPK3 expression. Stage 2 cells are enlarged with loss of CD20^+^ cytoplasm and increased expression of RIPK3 associated with CD20^+^RIPK3^+^ nodal latticework. In stage 3, only largely emptied cell “ghosts” outlined by residual staining of latticework nodes are visible. The progressive emptying of the cells is mediated through the nodal latticework illustrated in the exploded view of the PANoptosome (Fig. 18). Nuclear changes typical of apoptosis-heterochromatin condensation, nuclear bodies and marginalized envelope, provide visual evidence of B cell PANoptosis, in this case, necroptosis + apoptosis.

**Fig. 16.**
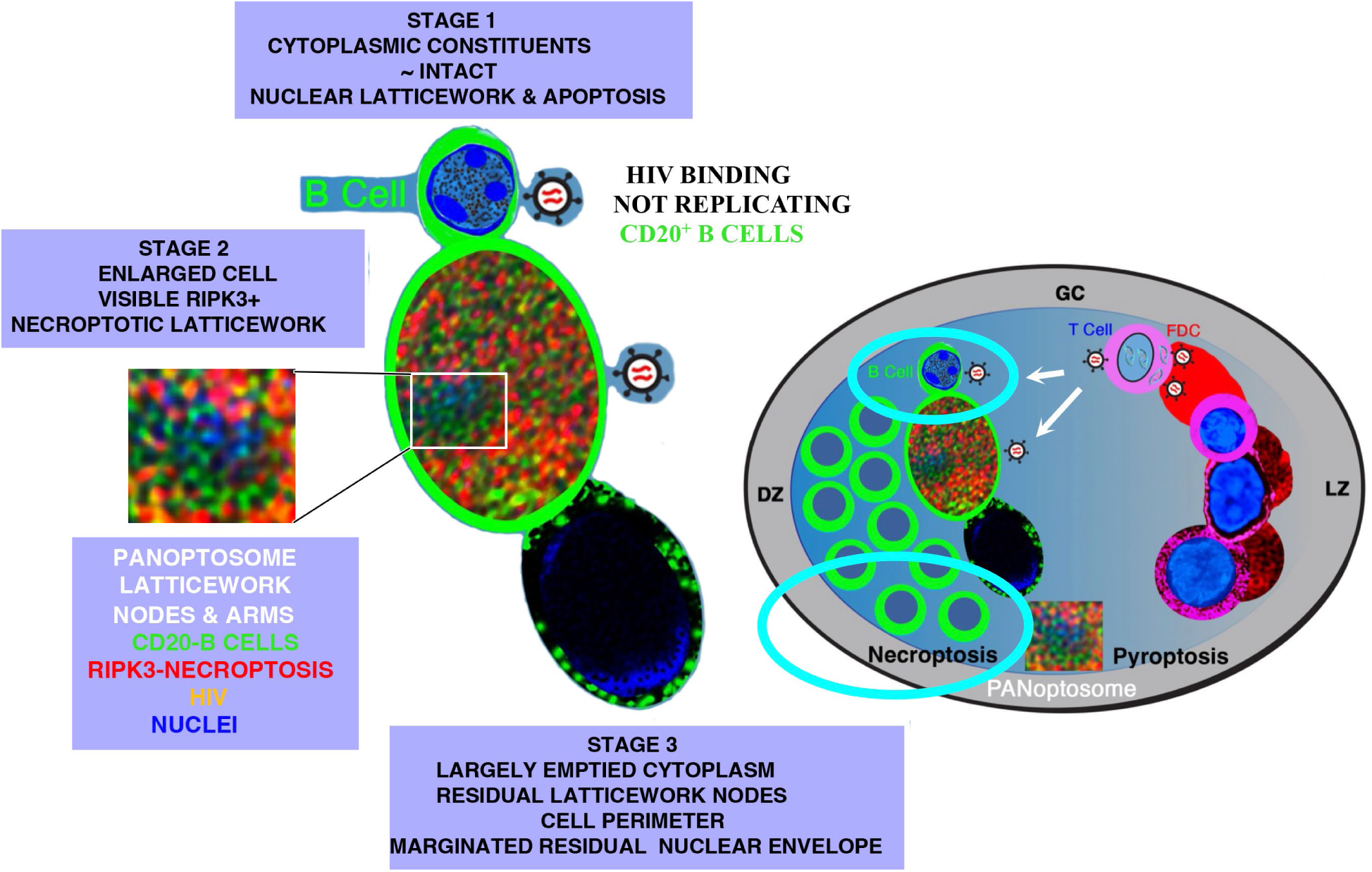
B cell necroptosis graphic. HIV produced by T cells bound to B cells without replication (intracellular-HIV RNA-negative) is associated with necroptosis of the B cells. Necroptosis proceeds through three stages illustrated and defined in the graphic. Illustrations of nuclear changes, latticework and PANoptosome nodes and arms (exploded view connected to the stage 2 cell) are actual images from the tissue samples.

**Fig. 17.**
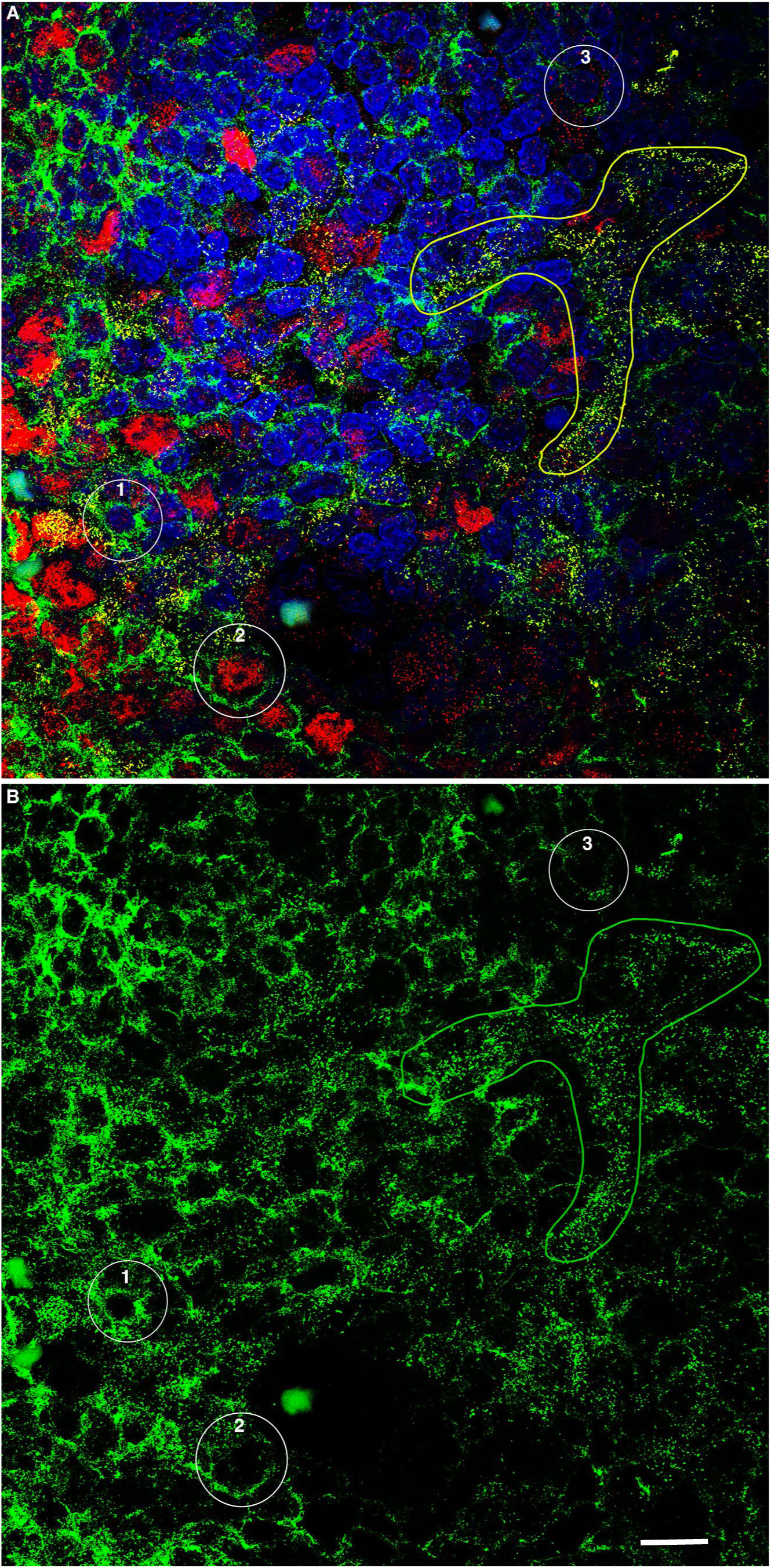
HIV binding and associated necroptosis in a B cell syncytium. NSPARC super-resolution images. HIV virions, yellow; CD20, green; RIPK3, red; DAPI, blue. **A.** HIV + CD20 + RIPK3 + DAPI. **B.** CD20. **A, B.** HIV bound to a syncytium of fused CD20^+^ B cells. Yellow line in **A** traces an example of HIV bound to CD20 (green line in **B**). Note the lack of discernable intracellular HIV RNA in the B cells as evidence of virus binding without replication. The numbered white circles identify examples of cells in early through later stages of necroptosis. Stage 1 cells are small with a largely intact rim of CD20^+^ cytoplasm and low levels of RIPK3 expression. Stage 2 cells are enlarged with progressive emptying of CD20^+^ cytoplasm surrounding RIPK3^+^ nodal latticework. In stage 3, residual CD20^+^RIPK3^+^ latticework nodes surround largely empty cells. Scale bar = 10 microns.

**Fig. 18.**
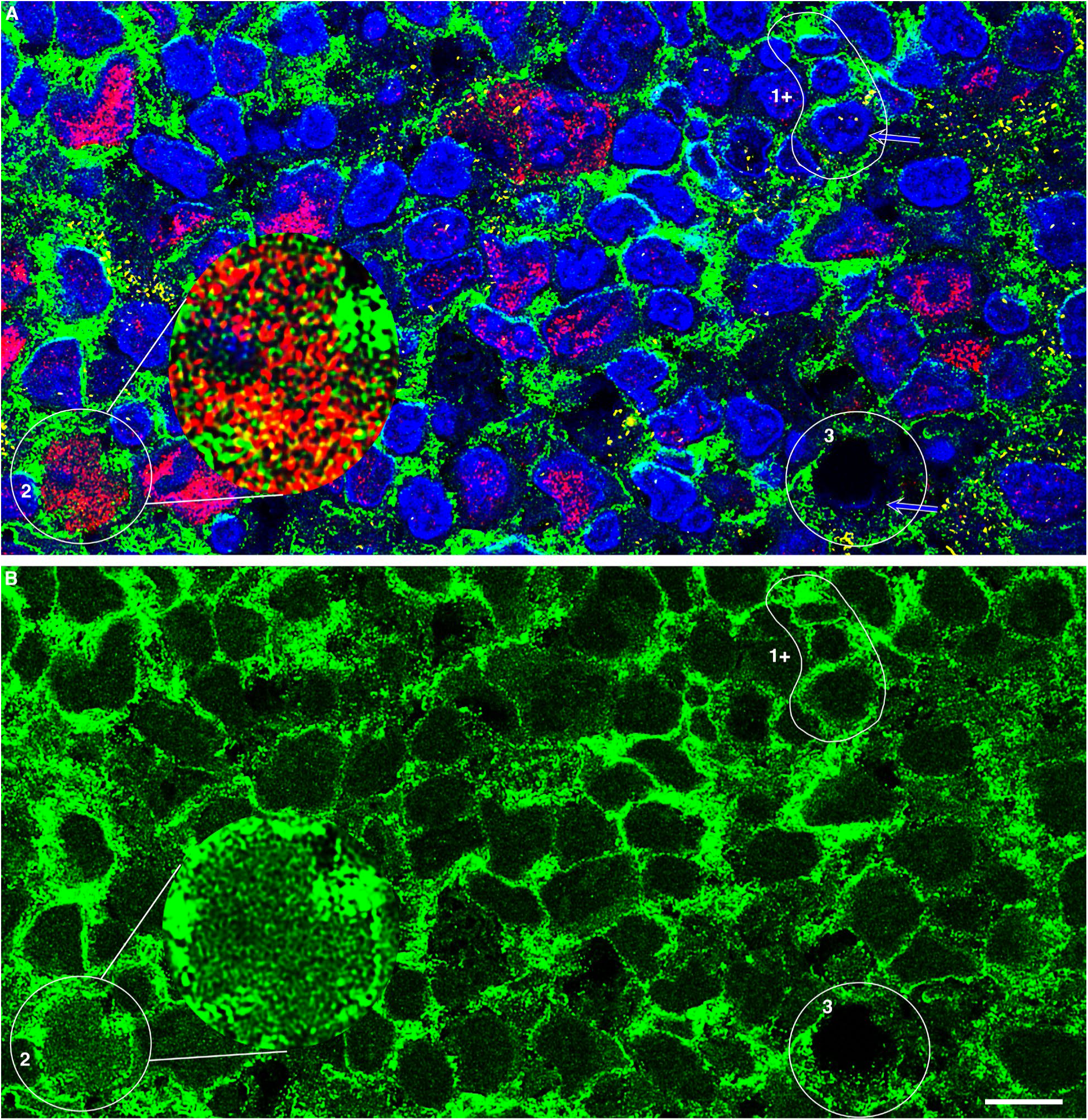
Progressive stages of necroptosis of CD20^+^ B cells and PANoptosome nodal latticework. NSPARC super-resolution images. HIV virions, yellow; CD20, green; RIPK3, red; DAPI, blue. **A.** HIV + CD20 + RIPK3 + DAPI. **B.** CD20. **A.** Numbered traces illustrate progressive stages in B cell necroptosis: Stage 1+: White trace outlines progressive enlargement of three fused B cells and nuclei; emptying of cytoplasm and rim comprised of CD20^+^ nodes; minimal expression of RIPK3. White outlined blue arrow points to apoptotic appearing nucleus with marginated envelope and nuclear fragments. Stage 2. Enlarged cells with thin rim of residual cytoplasm and CD20^+^ nodes, arms and latticework; increased expression of RIPK3; fragmented nucleus. Exploded view of the PANoptosome nodal latticework with red RIPK3^+^ nodes, green CD20^+^ nodes, HIV^+^ yellow nodes, and blue DAPI^+^ nodes. Note decreasing size of nodes associated with emptying of cellular and programmed cell death pathway contents. Stage 3. Necroptotic cells largely cleared of cytoplasm and RIPK3^-^. White outlined blue arrow points to residual visible nuclear envelope. **B.** CD20^+^ nodal latticework in stages 1-3 as described in **A**. Scale bar = 10 microns. Exploded view shows CD20^+^ PANoptosome nodal latticework.

### HIV-producing resting T cells persist in lymphoid tissues during ART when virus is undetectable in peripheral blood

By month 6 of ART, virus was undetectable in plasma, but HIV-producing cells were reduced in LN biopsies only about 80-fold and still easily detectable (Table 3; Fig. 19). The persisting HIV-producing cells were still CD4^+^CD25^-^ resting T cells (94 to 97 percent; Table 3). In contrast to D0 before ART, the majority of persisting cells with one exception were in the TZ (Table 3; Fig. 20) as neighbors or conjugates of two or more cells except for persistence of HIV-producing cells in GC, which recapitulated the before ART picture of massive fusion in the LZ (Fig. 20).

**Fig. 19.**
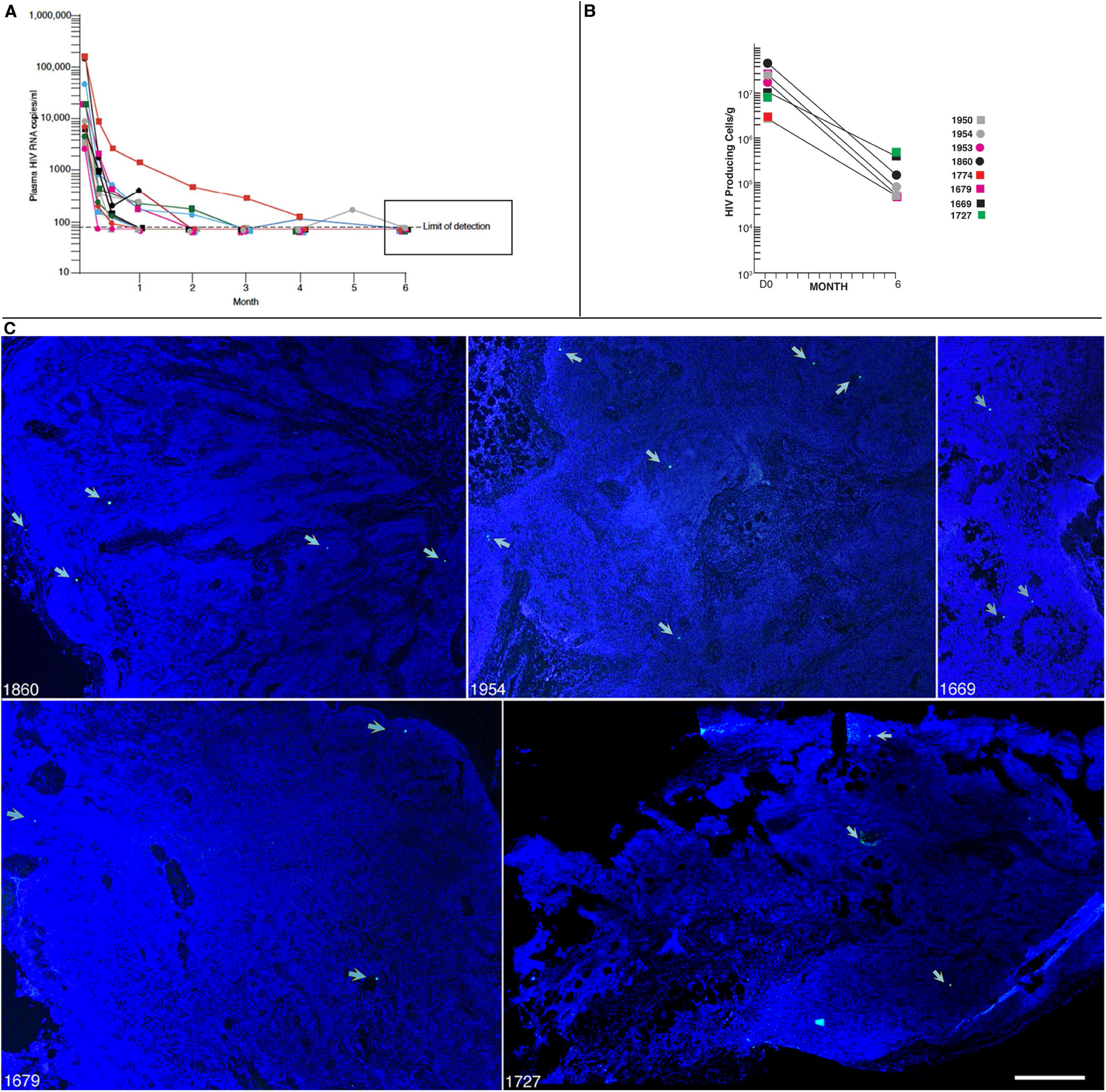
Persistence of HIV-producing cells in lymphoid tissues during ART. **A.** Viral decay in peripheral blood. Plasma viral loads reduced to the limits of detection at month 6 of ART. **B.** HIV-producing cells/g reduced about 80-fold in lymphoid tissues. **C.** 4X images of tissue sections from the PIDs shown with white outlined cyan arrows pointing to examples of single or two proximate HIV-producing cells (see Fig. 20) to illustrate ease of detection. Scale bar = 500 microns.

**Fig. 20.**
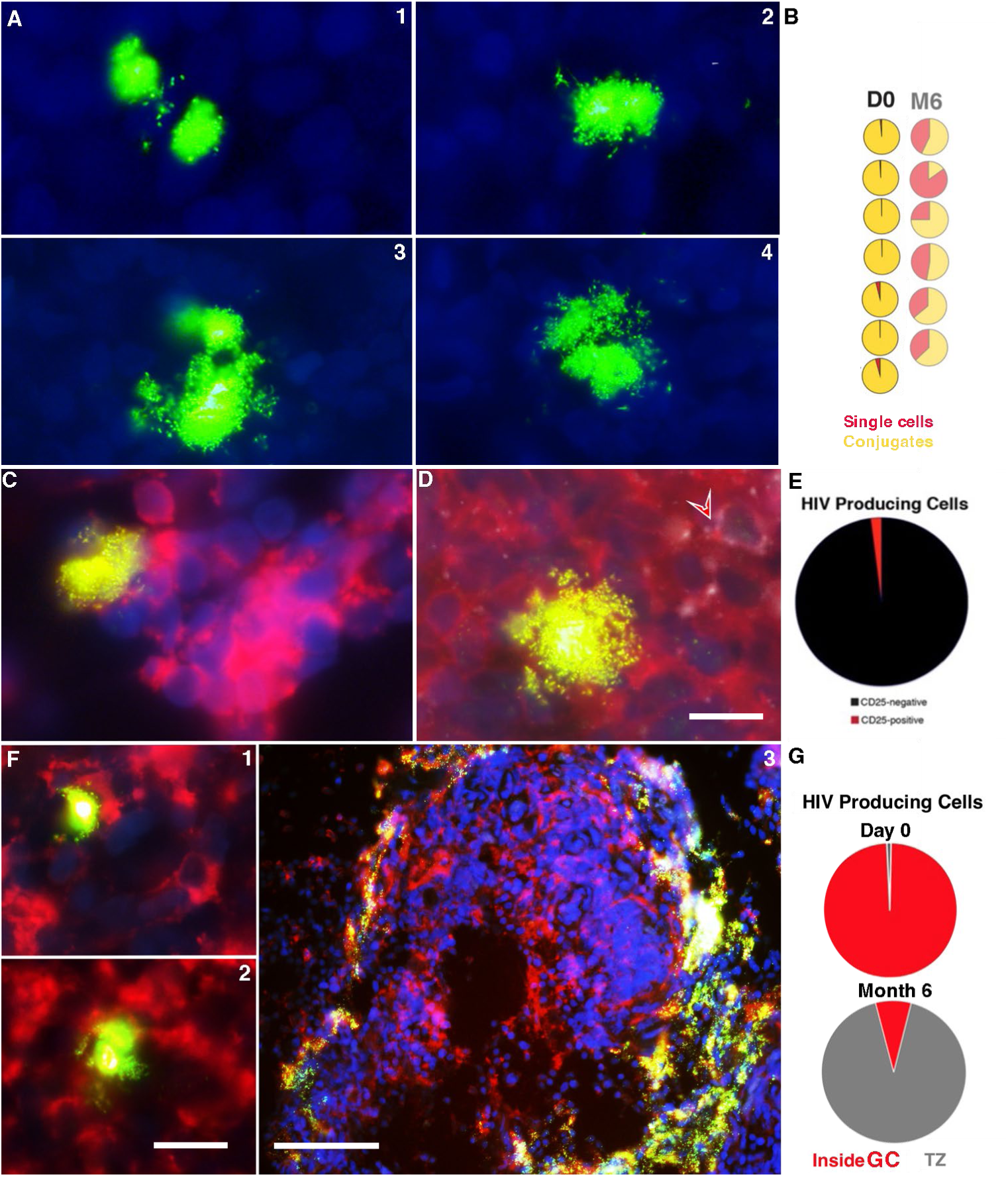
Persistence of HIV-producing CD4^+^CD25^-^ T cells in the TZ and GCs at month 6 of ART. **A.** 1-4. HIV virions and RNA, green. Examples of spatially proximate HIV-producing cells and cell conjugates. **B.** Pie sector diagrams of HIV-producing single cells and cell conjugates at D0 and M6 of ART showing increased proportion of single cells at M6 of ART. **C.** HIV virions and RNA (green) associated with red CD4^+^ cells are yellow. DAPI-stained nuclei are blue. **D.** HIV virions and RNA (green) associated with red CD4^+^CD25^-^ T cells are yellow. White outlined red arrow points to a HIV^-^CD4^+^CD25^+^ cell. Scale bar = 10 microns. **E.** Pie sector diagram showing the predominance (>90%) of CD4^+^CD25^-^ HIV producing T cells at M6 of ART. **F1-2**. Scale bar = 20 microns. Examples of yellow HIV producing CD4^+^ single cells and cell conjugates in a field of red CD4^+^ T cells in the TZ. **F3**. Scale bar = 100 microns. Rare M6 GC with fused and disrupted T cells fused to FDCs like D0. **G.** Pie sector diagrams showing the predominance of HIV-producing cells inside GCs at D0 and reversed pattern of HIV-producing T cells in the TZ at M6 of ART.

**Table 3.** HIV-producing/CD4^+^CD25^-^ T cells in GCs and TZ, Month 6 of ART.

| **PID** | **Total**<br>**HIV<sup>+</sup>**<br>**cells/g** | **Percent of**<br>**HIV<sup>+</sup> cells**<br>**in GCs** | **Percent of**<br>**HIV<sup>+</sup> cells**<br>**in TZ** | **Percent**<br>**HIV<sup>+</sup> cells**<br>**HIV<sup>+</sup>CD4<sup>+</sup>** | **Percent**<br>**HIV<sup>+</sup> cells**<br>**HIV<sup>+</sup>CD25<sup>-</sup>** |
| --- | --- | --- | --- | --- | --- |
| 1950 | 6.0 x 10<sup>4</sup> | 35.7 | 64.3 | 100 | 100 |
| 1954 | 8.2 x 10<sup>4</sup> | 16.6 | 83.4 | 83 | 96.3 |
| 1860 | 1.6 x 10<sup>5</sup> | 37.8 | 62.2 | 93 | 95.5 |
| 1679 | 9.1 x 10<sup>4</sup> | 13.6 | 86.4 | 88.6 | 100 |
| 1669 | 4.0 x10<sup>5</sup> | 71.6 | 28.4 | 98.7 | 92 |
| 1727 | 4.7 x 10<sup>5</sup> | 48.5 | 51.5 | 100 | 100 |
| **Averages** | **2.1 x 10<sup>5</sup>** | **37.3** | **62.7** | **93.9** | **97.3** |

**Quantification of HIV-producing cells.** HIV-producing cells in GCs and TZs in ~20 sections per patient were imaged under 40× magnification and enumerated manually and independently by two investigators and total HIV<sup>+</sup> cell counts normalized to tissue weight (grams). The proportion of HIV<sup>+</sup> cells located within GCs and TZ were calculated as percent of total HIV<sup>+</sup> cells.

## Discussion

### HIV-producing resting CD4^+^ T cells monopolize virus production and persistence

Here we show that HIV producing resting memory T cells in the lymphoid tissue reservoir in the chronic stage of infection account for nearly the entire population of productively infected cells before ART is initiated and the persistent productively infected population during ART. These studies complement those of the earliest detectable stage of HIV infection where we similarly showed that resting T cells dominate virus production and persistence in lymphoid tissues (LT) through two to five years of ART (10) and represent the latest chapter in over two plus decades on the critical role resting T cells play in transmission, propagation and persistence of infection in the LT reservoir in SIV and HIV infections (24–27). To put the sizes of the HIV-producing resting T cell populations in the HIV LT reservoir in perspective before and during ART, assuming that lymph nodes are representative of lymphoid tissues throughout the body, we estimate from the HIV-producing cells/g in the lymphoid tissue reservoir that a 70 kg person would contain about 200 billion HIV-producing cells before ART and about 150 million HIV-producing cells during ART (18).

### Why lymphoid tissue resting T cells dominate virus production and persistence

The susceptibility, availability, LT immunology, milieu and organization we think account for virus production in GCs by resting memory CD4^+^CD25^-^CD45RO^+^ T cells. First, susceptibility: the long-held concept that HIV is produced by activated T cells is based on in vitro experiments with cultured T cells (1) from peripheral blood (PB) that generally need to be activated to support virus production, particularly to induce pTEF-b, a critical transcription factor for HIV replication (28–31). However, CD4^+^ T cells in LT with a resting memory phenotype express pTEFb (10). Second, availability and contiguity: resting CD4^+^ T cells are essentially the only available spatially contiguous target cells to support cell-to-cell infection as shown previously in acute HIV infection (10) and here in the chronic stage of infection. Third, apposition of virus and target cells: the deposition of a replicating “antigen” on the FDCs creates an efficient high MOI infection complex of HIV-bearing FDCs underlying resting CD4^+^ T cell targets to concentrate multi-copy HIV DNA infection in the LZ. During ART, target cell availability and contiguity in GC and in the TZ continue to dictate productive infection of resting T cells in the chronic stage of infection.

### Role of fusion of cells in propagation and cytopathic effects of HIV infection

Fusion to form multinucleated giant cells and syncytia that undergo cytolysis *in vitro* (in tissue culture) has long been recognized as characteristic of the cytopathology of the lentiviruses from the prototype, visna virus (32), to HIV (33). By contrast, the massive fusion of T cells to form multicellular aggregates and syncytia *in vivo* in HIV-infected LT we document is without precedent in the literature (34), reflecting advances in the technology and microscopy for tissue image analysis. For HIV infection in LT, this massive fusion of T cells and the fusion of T cells to FDCs and a concentrated source of virus serves as a mechanism for cell-to-cell propagation of infection and efficient production in a high MOI infection complex.

### PANoptotosomal destruction of GC populations and impaired humoral immunity

We found that fusion and syncytium formation are general features of the cytopathic effects of HIV production in GCs that destroy T cells, FDCs and B cells through PANoptosis. PANoptosis was mediated by the previously described nodal latticework of the PANoptosome (23). We now present NSPARC super-resolution resolution images of the PANoptosome in which cell contents and PCD pathway components are concentrated in a nodal latticework structure that progressively empties and disrupts cells in stages of an asynchronous process that was visible in fixed tissues. We think it likely that the HIV envelope is not only responsible for the fusion of GC populations but also the initiator of PANoptosis, in HIV-producing T cells and from the virions produced bound by FDCs and B cells independent of replication, showing *in vivo* for B cells what had previously been found *in vitro* in B cells cultured from peripheral blood (35, 36). From an immunologically functional perspective, the PANoptotic destruction of the interacting T and B cell and FDC populations required for antibody production could certainly contribute to the well documented B cell abnormalities and impaired immune responses to HIV and vaccines (37–40).

### High multiplicity cell-to-cell infection and fusion in GCs and ART resistance

The nearly universal mode of cell-to-cell infection and fusion in the high MOI infection complexes in GCs raised the question of whether these complexes and mode of transmission would be resistant to ART, as has been suggested by *in vitro* models (41). We found to the contrary that these multicellular infectious complexes did not persist during ART. Rather, HIV-producing CD4^+^ T cells persisted in reduced numbers largely as single cells, doublets or cells in close proximity in the TZ and GC in varying proportions. We attribute the clearing of GCs of the high MOI complexes during ART to PANoptosis and to the reduced frequency of virus producing cells in a model that requires high levels of virus production for continuing formation and maintenance of the infection complexes.

### Persisting HIV-producing resting T cells and design and assessment of cure strategies

The challenge to curing HIV infections is currently generally thought to be recrudescent infection when treatment is interrupted from virus produced by reactivated latently infected cells established before ART was initiated (3–9). However, we have shown that HIV-producing resting CD4^+^ T cells persist after ART for two to five years initiated at the earliest stage of detectable infection (10) and we now show that HIV-producing resting CD4^+^ T cells persist during ART initiated in the chronic stages of infection when infection is usually diagnosed and treatment initiated. We emphasize the importance of LT analysis and the methodologies that support this conclusion and radical departure from the current view of the source of virus rebound. First, and most importantly, we think it unlikely that the HIV reservoir can be characterized and the source of virus rebound ascertained from studies of PB when virus-producing cells are in LT and not in the circulation. Second, even if LT samples were obtained, the PANoptotic destruction of HIV-producing resting T cells will have significantly reduced this population for analysis during the centrifugation steps to prepare samples and the parameters for single-cell analysis that would exclude the fused PANoptotic cell aggregates and debris. Third, the size of the replication-competent HIV reservoir is generally inferred by contemporary measures of the reservoir (42) rather than by the direct assessment we report here. We thus conclude that HIV cure strategies should be designed to incorporate assessment of HIV-producing resting T cells in LT and the impact of ART and new therapies by such high-resolution image analyses we describe that directly measure immediate sources of virus rebound, following an adage that *you can’t improve what you don’t measure*, revised here to *you can’t improve what you don’t see*. We further conclude that targeting the persistent HIV-producing resting T cell population should be a new focus and opportunity for HIV functional cure strategies.

## Methods

### Detecting and phenotyping HIV producing cells

HIV-producing cells were detected and phenotyped by RNAscope in situ hybridization and immunofluorescence. Formalin-fixed, paraffin-embedded (FFPE) 6 µm tissue sections were deparaffinized in xylene twice for 5 min each, followed by 100% ethanol twice for 5 min each. Sections were rehydrated in 80% ethanol for 5 min, rinsed in ddH₂O for 3 min, and post-fixed in freshly prepared 4% paraformaldehyde for 1 hour at room temperature. Slides were then washed in 80% ethanol twice for 5 min each, followed by dehydration in 100% ethanol for 5 min, and air-dried for 5 min. Sections were boiled for 10 minutes in RNAscope Target Retrieval Reagent (ACD, 322000), digested with Protease Plus 1:3 (ACD, 322330) at 40°C for 20 minutes, and then incubated at 40°C overnight with anti-sense probe V-HIV1-Clade B (ACD, 416111), with probe V-HIV-Clade AE (ACD, 446551) used as a negative control. Following hybridization, sections were washed in 0.5× RNAscope Wash Buffer, and signal amplification was performed using the RNAscope™ 2.5 HD Assay – RED Amplification Kit. Signals were developed using ELF97 (Invitrogen, E6588) at a 1:10 dilution at room temperature for approximately 20 minutes, with microscopic monitoring to achieve maximal signal intensity with no detectable background. For immunophenotyping, after RNAscope signal development, sections were washed twice in TBS for 5 minutes and blocked with Background Sniper (Biocare Medical, BS966) for 30 minutes at room temperature. For immunofluorescence staining, sections were incubated with primary antibodies at 4 °C overnight. Endogenous peroxidase activity was quenched with 3% hydrogen peroxide prior to signal amplification when Opal™ 570 were used. For single-antibody labeling, primary antibody was detected using Opal™ 570 or Alexa Fluor 555. For dual-antibody labeling, sections were first incubated with the initial primary antibody at 4 °C overnight, followed by Opal™ 570 development with sections incubated with 1X Opal Anti-Ms + Rb for 10 minutes at RT, washed in TBS for 5 minutes, and incubated with Opal™ 570 (1:100) for 10 minutes. After washing in TBS for 5 minutes, sections were blocked with Opal Antibody Diluent/Block buffer (Akoya, ARD1001EA) for 30 minutes and incubated with a second primary antibody (raised in a different species) at 4 °C overnight. Secondary antibodies conjugated to Alexa Fluor 488 were applied for 2 hours at room temperature. Alternatively, following the RNAscope protocol, two primary antibodies from different species were incubated simultaneously at 4 °C overnight, followed by species-specific Alexa Fluor 488 and 555 secondary antibodies applied for 2 hours at room temperature. After immunostaining, sections were washed in TBS, counterstained with DAPI, and mounted with Aqua-Polymount (Polysciences, 18606-20), and stored at 4 °C. Fluorescent images of HIV-producing cells were acquired using widefield fluorescence microscopy under WU illumination. Antibody staining images were captured using appropriate green and red filter sets at the same anatomical locations, the image sets were merged using Adobe Photoshop, and double- or triple-positive cells were enumerated manually.

### Detecting and phenotyping viral DNA in HIV producing cells

HIV DNA was detected by in situ hybridization, using our current pre-treatment protocol of a previously described method (43) and a HIV clade B sense probe (V-HIV1-CladeB-sense; ACD, 425531) to detect DNA, with the HIV clade AE sense probe (V-HIV-CladeAE-sense; ACD 478191) as a negative control. Following overnight hybridization at 40 °C, tissue sections were washed in 0.5×RNAscope wash buffer. For immunohistochemical detection, signal amplification was performed using the RNAscope™ 2.5 HD RED Amplification Kit. Chromogenic signal development was carried out with Warp Red Chromogen (1 drop of warp red chromogen mixed with 1.5 mL warp red buffer) for approximately 20 minutes. Sections were then washed twice in TBS for 5 minutes each, counterstained with hematoxylin for 1 minute, and rinsed in distilled water for 2 minutes. Slides were subsequently dehydrated by dipping 10 times in 100% ethanol, air-dried, cleared in xylene for 2 minutes, and cover slipped for imaging. Combined immunofluorescence detection of DNA and phenotyping was as described for HIV-producing cells, substituting the V-HIV1-CladeB-sense probe to detect DNA.

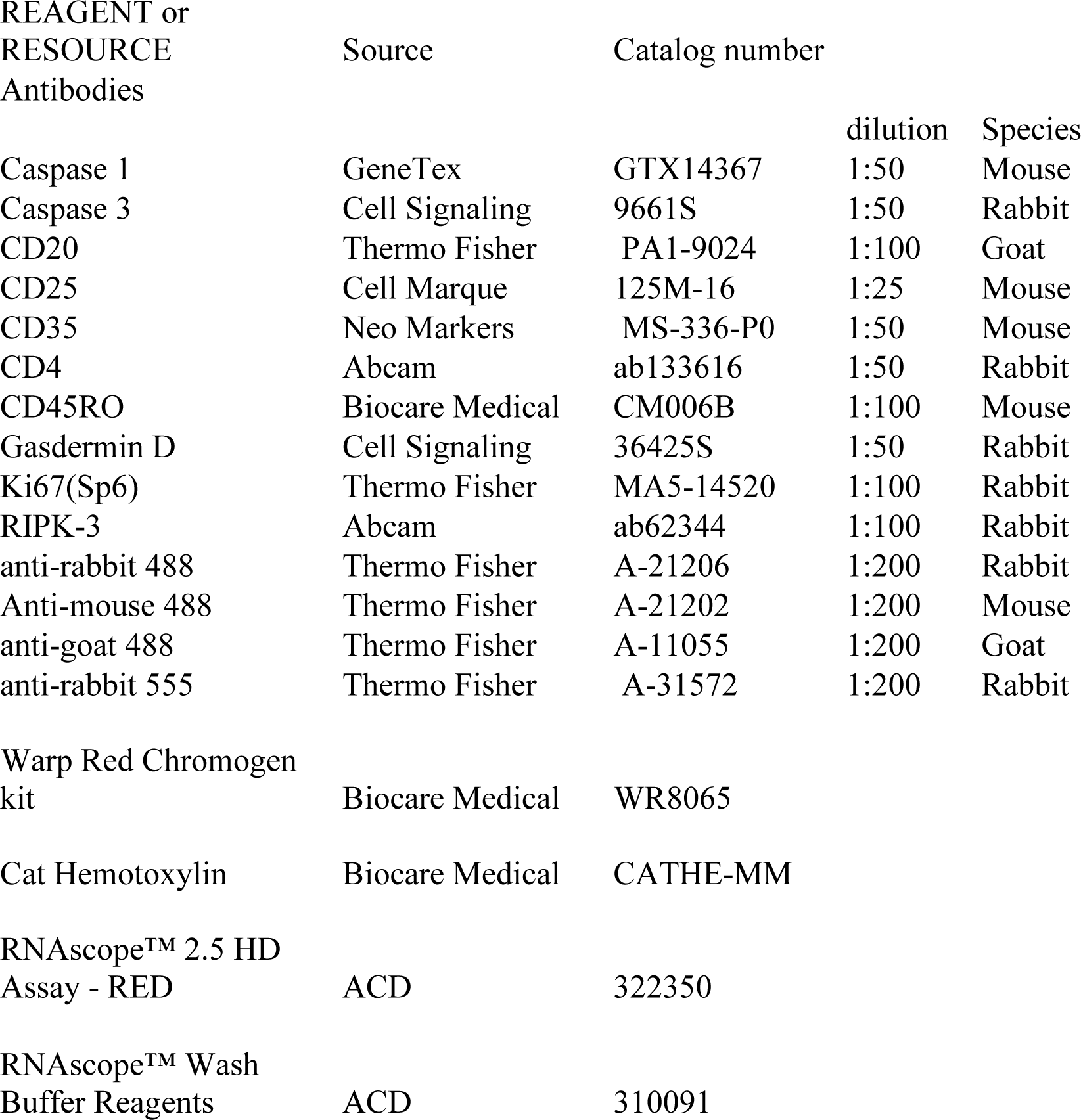
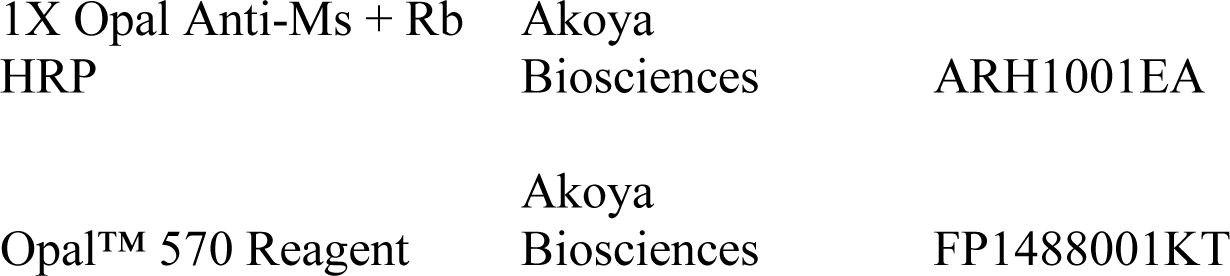

| REAGENT or<br>RESOURCE<br>Antibodies | Source | Catalog number | dilution | Species |
| --- | --- | --- | --- | --- |
| Caspase 1 | GeneTex | GTX14367 | 1:50 | Mouse |
| Caspase 3 | Cell Signaling | 9661S | 1:50 | Rabbit |
| CD20 | Thermo Fisher | PA1-9024 | 1:100 | Goat |
| CD25 | Cell Marque | 125M-16 | 1:25 | Mouse |
| CD35 | Neo Markers | MS-336-P0 | 1:50 | Mouse |
| CD4 | Abcam | ab133616 | 1:50 | Rabbit |
| CD45RO | Biocare Medical | CM006B | 1:100 | Mouse |
| Gasdermin D | Cell Signaling | 36425S | 1:50 | Rabbit |
| Ki67(Sp6) | Thermo Fisher | MA5-14520 | 1:100 | Rabbit |
| RIPK-3 | Abcam | ab62344 | 1:100 | Rabbit |
| anti-rabbit 488 | Thermo Fisher | A-21206 | 1:200 | Rabbit |
| Anti-mouse 488 | Thermo Fisher | A-21202 | 1:200 | Mouse |
| anti-goat 488 | Thermo Fisher | A-11055 | 1:200 | Goat |
| anti-rabbit 555 | Thermo Fisher | A-31572 | 1:200 | Rabbit |
| Warp Red Chromogen<br>kit | Biocare Medical | WR8065 |  |  |
| Cat Hemotoxylin | Biocare Medical | CATHE-MM |  |  |
| RNAscope™ 2.5 HD<br>Assay - RED | ACD | 322350 |  |  |
| RNAscope™ Wash<br>Buffer Reagents | ACD | 310091 |  |  |

|  |  |  |
| --- | --- | --- |
| 1X Opal Anti-Ms + Rb<br>HRP | Akoya<br>Biosciences | ARH1001EA |
| Opal™ 570 Reagent | Akoya<br>Biosciences | FP1488001KT |

### Confocal and NSPARC (Nikon Spatial Array Confocal) microscopy

Slide-mounted sections were examined on a Nikon AX R confocal Eclipse Ti microscope using the 20X CFI Plan Apochromat Lambda D 20X, 0.8 NA; CFI Plan Apochromat Lambda D 60X. 1.42 NA or CFI Plan Apochromat Lambda D 100X, 1.45 NA oil objective. Excitation laser wavelengths of 405 nm, 488 nm, 514 nm, and 561 nm were utilized sequentially. NSPARC confocal imaging was performed using SR mode using the optimal scan area zoom for each collection. All data were acquired in Galvo mode with confocal imaging using a pinhole size of 1.2 AU. Post acquisition duplicated images were processed using Nikon Elements V 6.02 using Deconvolution in NSPARC mode and look up tables adjusted for presentation.

## Acknowledgements

We thank Colleen O’Neill and Tim Leonard for preparation of the manuscript and figures; Peter Crawford, Michael Gale, Jr. and Harris Haase for helpful discussion; Jakub Tolar for discussion and support; Vivian Gao and Vincent Gao for technical help; and the resources and staff at the University of Minnesota University Imaging Centers (UIC). SCR_020997.

Research funding: National Institutes of Health R01 AI134406; Platou Medical Discovery Fund and Ashley T. Haase’s Regents’ Professorship, University of Minnesota.

## Declaration of interests

The authors declare no competing interests.

## Notes

### Competing Interest Statement

The authors have declared no competing interest.

## References

1. Folks, T. et al. Susceptibility of normal human lymphocytes to infection with HTLV-III/L. J. Immunol. 136, 4049–4053 (1986).

2. Ho, D. et al. Rapid turnover of plasma virions and CD4 lymphocytes in HIV-1 infection. Nature. 373, 123–126 (1995).

3. Siliciano, R.F. & Greene, W.C. HIV latency. Cold Spring Harb Perspect Med. 1, a007096 (2011).

4. Shan, L. et al. Transcriptional programming during effector-to-memory transition renders CD4^+^ T cells permissive for latent HIV-1 infection. Immunity 47, 766–775 (2017).

5. Chun, T.W. et al. Quantification of latent tissue reservoirs and total body viral load in HIV-1 infection. Nature 387, 183–188 (1997).

6. Finzi, D. et al. Identification of a reservoir for HIV-1 in patients on highly active antiretroviral therapy. Science 278, 1295–1300 (1997).

7. Wong, J. et al. Recovery of replication-competent HIV despite prolonged suppression of plasma viremia. Science 278, 1291–1295 (1997).

8. Margolis, D.M. et al. Curing HIV: seeking to target and clear persistent infection. Cell 181(1), 189–206 (2020).

9. Cohn, L.B., Chomont, N. & Deeks, S.G. The biology of the HIV-1 latent reservoir and implications for cure strategies. Cell Host Microbe. 27(4), 519–530 (2020).

10. Wietgrefe, S.W. et al. Initial productive and latent HIV infections originate in vivo by infection of resting T cells. J. Clin. Invest. 133(22), e171501 (2023).

11. Fletcher, C.V., et al. Persistent HIV-1 replication is associated with lower antiretroviral drug concentrations in lymphatic tissues. Proc. Natl. Acad. Sci. USA. 111(6): 2307-2312 (2014).

12. Wietgrefe S.W. et al. Detecting sources of immune activation and viral rebound in HIV infection. J. Virol. 96(15), e0088522 (2022).

13. Sakuragi, J. Morphogenesis of the infectious HIV-1 virion. Front. Microbiol. DOI=10.3389/fmicb.2011.00242 (2011).

14. Tenner-Racz, K. & Racz, P. Follicular dendritic cells initiate and maintain infection of the germinal centers by human immunodeficiency virus. Curr. Top. Microbiol. Immunol. 201, 141–159 (1995).

15. Fox, C.H. et al. Lymphoid germinal centers are reservoirs of human immunodeficiency virus type 1 RNA. J. Infect. Dis. 164, 1051–1057 (1991).

16. Heath, S.L., Tew, J.G., Szakal, A.K. & Burton, G.F. Follicular dendritic cells and human immunodeficiency virus infectivity. Nature 377, 740–744 (1995).

17. Burton, G.F. et al. Follicular dendritic cell contributions to HIV pathogenesis. Semin. Immunol. 4, 275–84 (2002).

18. Haase, A.T. et al. Quantitative image analysis of HIV-1 infection in lymphoid tissue. Science 274, 985–989 (1996).

19. Cavert, W. et al. Kinetics of response in lymphoid tissues to antiretroviral therapy of HIV-1 infection. Science 276, 960–964 (1997).

20. Haase, A.T. Population biology of HIV-1 infection: Viral and CD4+ T cell demographics and dynamics in lymphatic tissues. Ann. Rev. Immunol. 17, 625–656 (1999).

21. Christgen, S. et al. Identification of the PANoptosome: a molecular platform triggering pyroptosis, apoptosis, and necroptosis (PANoptosis). Front. Cell. Infect. Microbiol. 10, 237 (2020).

22. Samir, P., Malireddi, R.K.S. & Kanneganti, T.D. The PANoptosome: A deadly protein complex driving pyroptosis, apoptosis, and necroptosis (PANoptosis). Front. Cell. Infect. Microbiol. 10, 238 (2020).

23. Schifanella, L. et al. The defenders of the alveolus succumb in COVID-19 pneumonia to SARS-CoV-2 and necroptosis, pyroptosis, and PANoptosis, J. Infect. Dis. 227, jiad056, 10.1093/infdis/jiad056 (2023).

24. Zhang, Z.-Q. et al. Sexual transmission and propagation of simian and human immunodeficiency viruses in two distinguishable populations of CD4^+^ T cells. Science 286, 1353–1357 (1999).

25. Zhang, Z.-Q., et al. Roles of substrate availability and infection of resting and activated CD4+ T cells in transmission and acute simian immunodeficiency virus infection. Proc. Natl. Acad. Sci. USA. 101, 5640-5645 (2004).

26. Li, Q. et al. Peak SIV replication in “resting” memory CD4^+^ T cells depletes gut lamina propria CD4^+^ T cells. Nature 434, 1148–1152 (2005).

27. Reilly, C., Wietgrefe, S., Sedgewick, G. & Haase, A.T. Determination of simian immunodeficiency virus production by infected activated and resting cells. AIDS 21, 163–168 (2007).

28. Pan, X. et al. Restrictions to HIV-1 replication in resting CD4^+^ T lymphocytes. Cell Res. 23, 876–885 (2013).

29. Ramakrishnan, R. et al. Making a short story long: regulation of P-TEFb and HIV-1 transcriptional elongation in CD4^+^ T lymphocytes and macrophages. Biology (Basel) 1, 94–115 (2012).

30. Tyagi, M., Pearson, R.J. & Karn, J. Establishment of HIV latency in primary CD4^+^ cells is due to epigenetic transcriptional silencing and P-TEFb restriction. J. Virol. 84, 6425–6437 (2010).

31. Karn, K. The molecular biology of HIV latency: Breaking and restoring the Tat-dependent transcriptional circuit. Curr. Opin. HIV AIDS. 6, 4–11 (2011).

32. Haase, A.T. The slow infection caused by visna virus. Curr. Top. Microbiol. Immunol. 72, 101–156 (1975).

33. Lifson, J.D. et al. AIDS retrovirus induced cytopathology: giant cell formation and involvement of CD4 antigen. Science. 232, 11231127 (1986).

34. Compton, A.A. & Schwartz, O. They might be giants: does syncytium formation sink or spread HIV infection? PLoS Pathog. 13(2), e1006099 (2017).

35. Embretson, J. et al. Massive covert infection of helper T lymphocytes and macrophages by HIV during the incubation period of AIDS. Nature 362, 359–62 (1993).

36. Moir, S. et al. B cells of HIV-1-infected patients bind virions through CD21-complement interactions and transmit infectious virus to activated T cells. J. Exp. Med. 192, 637–46 (2000).

37. Moir, S. & Fauci, A.S. B-cell responses to HIV infection. Immunol. Rev. 275, 33–48 (2017).

38. Ballet, J.J. et al. Impaired anti-pneumococcal antibody response in patients with AIDS-related persistent generalized lymphadenopathy. Clin. Exp. Immunol. 68, 479–487 (1987).

39. Miotti, P.G. et al. The influence of HIV infection on antibody responses to a two-dose regimen of influenza vaccine. J. Amer. Med. Assoc. 262, 779–783 (1989).

40. Opravil, M., Fierz, W., Matter, L., Blaser, J. & Luthy, R. Poor antibody response after tetanus and pneumococcal vaccination in immunocompromised, HIV-infected patients. Clin. Exp. Immunol. 84, 185–189 (1991).

41. Sigal A, et al. Cell-to-cell spread of HIV permits ongoing replication despite antiretroviral therapy. Nature 477, 95–98. (2011).

42. Abdel-Mohsen, M. et al. Recommendations for measuring HIV reservoir size in cure-directed clinical trials. Nat. Med. 26, 1339–1350 (2020).

43. Deleage, C., et al. Defining HIV and SIV reservoirs in lymphoid tissues. Pathog. Immun. 1, 68–106. (2016).

