## Supplemental Fig 1 Legend for "HIV-producing resting T cells monopolize virus production, associated PANoptosis, and persistence in lymphoid tissue"

### Supplemental Information

**Supplemental Fig. 1. Movie of HIV RNA$^+$GSD$^+$ pyroptotic nodes in fused CD35$^+$ FDCs.** HIV, yellow; CD35$^+$, green; GSD, red; nuclei, blue. Movie shows a 3D view from **Fig. 14** of viral and pyroptotic GSD contents of latticework nodes in the fused FDCs. Stepwise examination at 200% of z-slice sequences from 00.33 through the end of 00.48 seconds in the movie reveals the nodal latticework in the FDCs and nuclei, coincidence of HIV, GSD and CD35 in the nodes, and asynchronous progression of clearance of nodal contents that places the smaller nodes at the most advanced stage of clearance compared to the larger nodes at the earlier stages.
